# Investigating the dynamics of microbial consortia in spatially structured environments

**DOI:** 10.1101/2020.02.17.953240

**Authors:** Sonali Gupta, Tyler D. Ross, Marcella M. Gomez, Job L. Grant, Philip A. Romero, Ophelia S. Venturelli

## Abstract

The spatial organization of microbial communities arises from a complex interplay of biotic and abiotic interactions and is a major determinant of ecosystem functions. We design a microfluidic platform to investigate how the spatial arrangement of microbes impacts gene expression and growth. We elucidate key biochemical parameters that dictate the mapping between spatial positioning and gene expression patterns. We show that distance can establish a low-pass filter to periodic inputs, and can enhance the fidelity of information processing. Positive and negative feedback can play disparate roles in the synchronization and robustness of a genetic oscillator distributed between two strains to spatial separation. Quantification of growth and metabolite release in an amino-acid auxotroph community demonstrates that the interaction network and stability of the community are highly sensitive to temporal perturbations and spatial arrangements. In sum, our microfluidic platform can quantify spatiotemporal parameters influencing diffusion-mediated interactions in microbial consortia.

## INTRODUCTION

Spatial organization is a prevalent feature of microbiomes ranging from soil^1^ to the human gastrointestinal tract^2^. This spatial structure is a major driver of microbial community behaviors, stability, and responses to environmental perturbations^3–7^. The spatial organization of microbiomes span multiple scales: variation in environmental (abiotic) parameters dictate behaviors over longer length scales (centimeter to meter), whereas inter-microbial interactions impact community behaviors on shorter length scales (tens of microns to centimeters)^8,9^. It is not currently understood how a microbiome’s spatial structure impacts community function and stability.

The proximity of members within a community is a major determinant of the costs and benefits of microbial interactions, and shapes how the ecological network evolves over time^10,11^. Spatial structure is known to provide ecological benefits, such as promoting population survival through local public good production^12^ and enhancing biofilm resilience to environmental perturbations^5,13^. Spatial heterogeneity was shown to reduce the propensity for a prisoner’s dilemma by enabling the stable coexistence of cooperator and cheater populations^3^. Ecosystem properties are shaped by the balance between cooperation and competition, which have been shown to dominate in different spatial regimes^14^. The degree of mixing of strains impacts the concentrations of resources and toxins in local microenvironments, which in turn dictates the outcome of invasion of non-resident organisms into the community^15,16^.

The majority of microbial interactions are mediated by diffusible compounds including metabolites, chemical signals, or proteins^7^. These biomolecular mediators can enhance or inhibit community member’s growth rates, as well as modify the activities of intracellular signaling or metabolic networks. In spatially heterogenous environments, such as the human gut^2^ or plant rhizosphere^17^, community-level functions are driven by factors such as physical separation of bacterial populations, bulk flow of biomolecules, and cell motility. In the absence of convection and cell motility, amino-acid auxotrophs were shown to interact over tens of microns^18^, demonstrating that metabolite secretion impacts groups of cells occupying a small local neighborhood. However, the spatial interaction range for quorum-sensing mediated bacterial communication was expanded by amplifying the production of chemical signals via positive feedback^19^.

It is challenging to study and control spatial arrangements of bacterial populations on the micron scale, a regime where microbial interactions can significantly impact spatial distributions^8,9^. Microfabrication of patterned agarose^20^, hydrogels^21^, partitioned microfluidics^22^, nanoporous membranes^14^, cellulose nanofibrils^23^, nanochannels^24^ and bioprinting^25^ techniques have been used to physically separate bacterial populations to study interactions and chemical signal communication. A platform that integrates micron-level spatial patterning, temporal control of environmental stimuli, and single-cell quantification of cell growth and gene expression has not been fully leveraged to study diffusion-mediated interactions in microbial communities. A detailed and quantitative understanding of how spatiotemporal parameters impact microbial communities would provide insights into the functional roles of spatial organization and how spatial structure could be manipulated for biomedical or biotechnological applications.

We develop a microfluidic platform, MISTiC (Mapping Interactions across Space and Time in Communities), to interrogate spatiotemporal parameters in microbial consortia. MISTiC enables temporal control of environmental inputs, precise spatial positioning of bacterial populations on the micron-scale, and long-term time-lapse imaging of growth and gene expression in continuous culture at the single-cell and population levels. We study how spatial separation impacts unidirectional quorum-sensing communication in a synthetic *E. coli* sender-receiver consortium. A dynamic computational model elucidates key biochemical parameters that modulate information transmission between the sender and receiver across distance. Using this system, we demonstrate that distance can enhance the fidelity of information transmission in specific input frequency regimes. To investigate the principles of systems shaped by feedback, we examine the dynamics of an engineered *E. coli* consortium that exhibits coupled gene expression oscillations via inter-strain bidirectional communication and dual-feedback loops. We show that each strain exhibited substantially different response to changes in the interaction length parameter, suggesting that positive and negative feedbacks play opposing roles in shaping the robustness of oscillations to spatial separation.

To explore the effects of distance on metabolic interactions, we study the growth dynamics and metabolite release of an *E. coli* amino acid auxotroph community at the single-cell and population-levels. Our results show an asymmetry in the interaction network, which can be further enhanced by spatial separation of tens of microns. Metabolite measurements demonstrate that the strains exhibit disparate amino acid release profiles as a function of growth and amino acid availability, providing insights into the interaction network and community stability. Together, these data show that MISTiC can be used to precisely quantify the role of micron-scale spatial separation and temporal perturbations on microbial interactions, community functions and stability.

## RESULTS

### A microfluidic platform to interrogate microbial interactions

Previous studies have demonstrated that pairwise interactions drive the behaviors of complex multi-member communities^26–28^. Therefore, we designed MISTiC to probe diffusion-mediated interactions between pairs of strains by separating bacterial populations into growth chambers that are connected by interaction channels (**Fig. 1a**). Our microfluidic design balances the pressures between interacting growth chambers, and thus prevents convective flow through the interaction channel^29^. This allows us to directly study the effect of molecular diffusion between interacting cell populations. Each 10 × 50 × 1 μm growth chamber contains approximately 150 cells that are restricted to a monolayer for real-time quantification of gene expression and growth. The interaction channels are less than 0.5 μm tall and structurally supported by 0.5 μm pillars, which serves as a physical barrier for the cells, while permitting diffusion of biomolecules between growth chambers. Our MISTiC device contains 25, 50, 100, and 250 μm interaction channels to study how distance on this scale influences microbial interactions. As the population grows and divides, excess cells are washed away by continuous media flow, enabling long-term imaging.

**Figure 1.**
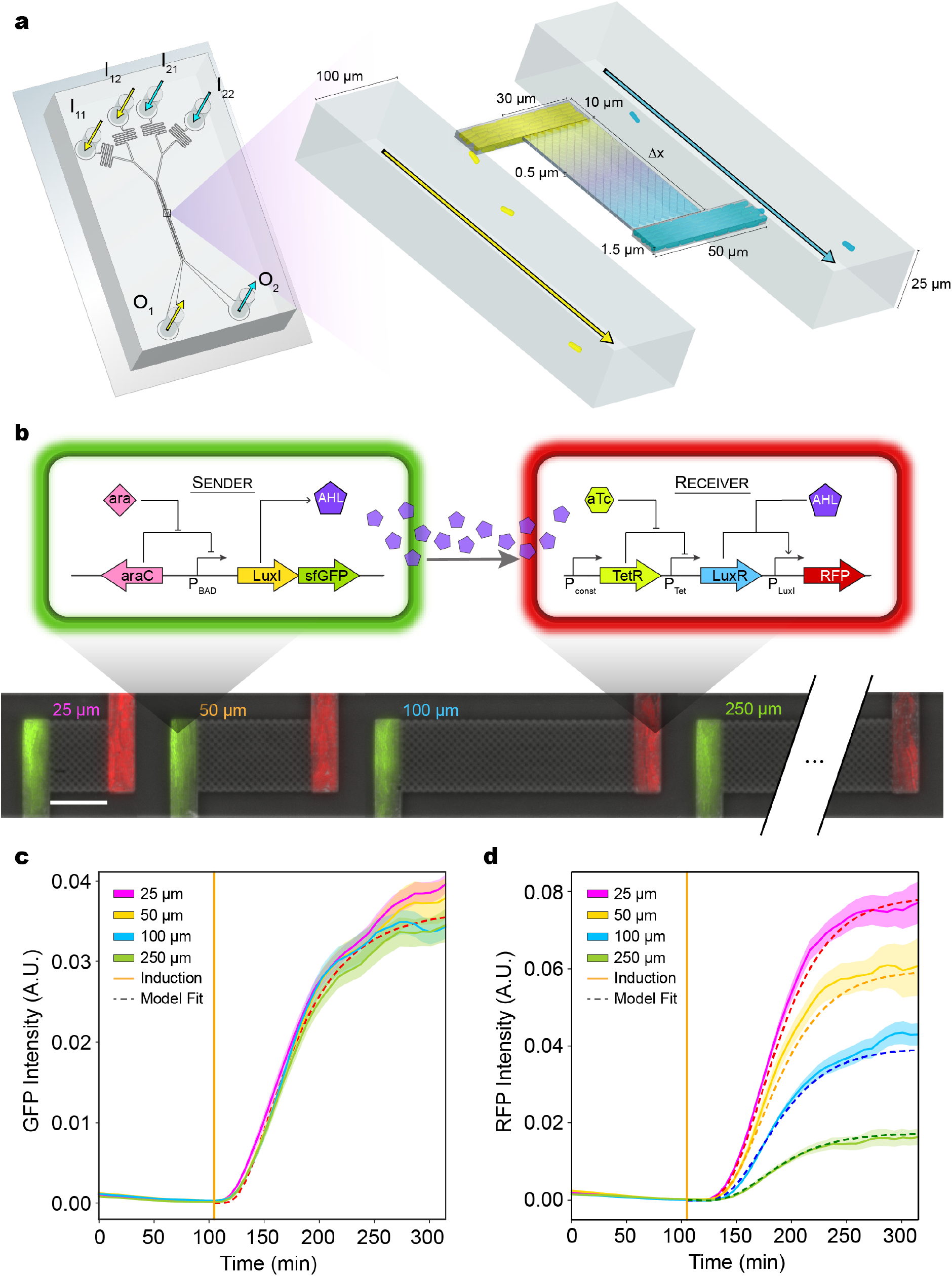
Design of a microfluidic platform, MISTiC, to investigate the role of spatiotemporal parameters on inter-strain communication. **(a)** Schematic of the microfluidic device. The inlets I_11_, I_12_ or I_21,_ I_22_ connect to the outlets O_1_ or O_2_, respectively, and allow temporal control of the environmental conditions for each strain. Cells are initially seeded into growth chambers and grow to reach a maximum population size of approximately 150 cells. Continuous flow of media through the main channels removes excess cells. Pairs of growth chambers are separated by a lattice of pillars, defined as the interaction channel, which allows diffusion of biomolecules and prevents cells from entering the interaction channel. The device has ten pairs of growth chambers for each separation distance. **(b)** Top: Schematic of the genetic circuit in the sender and receiver *E. coli* strains. In the sender strain, the operon containing the synthetase LuxI and GFP is induced in response to arabinose. The synthetase LuxI produces the acyl-homoserine lactone (3-oxo-C6-HSL or AHL), which diffuses through the interaction channel into the receiver strain growth chamber. In the receiver strain, AHL binds to LuxR to form an activated LuxR-AHL complex, which in turn activates RFP expression. Bottom: Overlaid representative fluorescence and phase contrast microscope images of the sender and receiver strains in the device for each interaction channel length (Experiment 1, **Table 1**). The scale bar represents 25 μm. **(c)** GFP fluorescence in sender growth chambers as a function of time. The vertical line indicates the time at which arabinose was introduced. Shaded regions represent one standard deviation from the mean. The dashed line denotes the model fit. **(d)** RFP fluorescence over time in the receiver growth chambers. The vertical line indicates the time at which arabinose was introduced. Shaded regions represent one standard deviation from the mean. The dashed line denotes the model fit. Source data are provided as a Source Data file.

**Table 1.** MISTiC experimental conditions. Pre-culture conditions refer to the environment within the microfluidic device from the beginning of the experiment to the time of the first media switch. Test culture conditions refer to the media conditions following the first media switch. Quorum sensing experiments were performed in Luria Broth (LB) whereas remaining experiments used MOPS EZ Rich Defined Medium, with the modifications specified above. AA refers to EZ amino acid solutions containing all amino acids and AA* refers to an amino acid solution lacking methionine (M) or phenylalanine (F). In experiments where F and M were added separately, 1X F and M refers to 0.4 mM and 0.2 mM, respectively. The pre-culture amino acid fraction was varied to control cell growth for allowing sufficient fluorescent reporter expression. Outliers refer to the number of paired growth chambers excluded for each distance (25 μm;50 μm;100 μm;250 μm) based on a set of specific criteria (**Materials and Methods**). ^**+**^For these experiments, the numbers represent single outlier growth chambers (20 total for each distance, 25 μm;50 μm;100 μm;250 μm) that were excluded from the analysis (**Materials and Methods**).

| Experiment | Descriptor | Media switch (min) | Pre-culture condition | Test condition | Outliers |
| --- | --- | --- | --- | --- | --- |
| 1 | Q.S. (step response) | 105 | aTc | aTc + ara | 1;1;0;0 |
| 2 | Q.S. (forced oscillator, 2 hr) | 140 | aTc | aTc ± ara | 1;1;1;5 |
| 3 | Q.S. (forced oscillator, 1 hr) | 184 | aTc | aTc ± ara | 0;0;1;3 |
| 4 | Dual-feedback oscillators | 217 | NA | IPTG | 0;0;3;1 |
| 5 | Auxotroph (control) | 218 | 0.25X AA + IPTG | 0.25X AA | 0;0;1;2 |
| 6 | Auxotroph (coupled) | 420 | 1X AA* + 0.1X F/M + IPTG | 1X AA* | 3;2;0;1 |
| 7 | Auxotroph (mixed) | 60 | 1X AA* + 0.1X F/M + IPTG | 1X AA* + IPTG | 5;2;2;1<sup>+</sup> |
| 8 | Auxotroph ($\Delta metA$ control) | 110 | 1X AA* + 0.1X F/M + IPTG | 1X AA* + IPTG | 1;2;3;8<sup>+</sup> |
| 9 | Auxotroph ($\Delta pheA$ control) | 90 | 1X AA* + 0.1X F/M + IPTG | 1X AA* + IPTG | 3;2;2;7<sup>+</sup> |
| 10 | Auxotroph (rescue - $\Delta met$) | 24 | 0.5X AA* + 1X M + | 0.5X AA* + 1X M | 0;0;0;1 |
|  |  |  | 0.05 X F<br>+ IPTG |  |  |
| 11 | Auxotroph<br>(coupled) | 710 | 0.1X AA<br>+ IPTG | 0.003X AA | 1;0;0;0 |
| 12 | Auxotroph<br>(coupled) | 148 | 0.1X AA<br>+ IPTG | 0.005X AA | 1;0;0;3 |
| 13 | Auxotroph<br>(coupled) | 285 | 0.2X AA<br>+ IPTG | 0.02X AA | 1;2;0;0 |

The environmental conditions can be dynamically controlled for each strain in the community using separate inlets. We characterized the molecular gradients established across interaction channels using a fluorescent dye (**Fig. S1**). A fixed concentration of fluorescein was loaded into the “source” chambers, which diffused across the interaction channel into the “sink” chambers. The average fluorescein concentrations within sink chambers decreased as a function of spatial separation between the source and sink chambers. The concentration of fluorescein within the interaction channels decreased as a function of distance from the source chamber. To capture diffusion across the interaction channels, we built a computational model that represented diffusion as a one-dimensional process by discretizing space into one-micron regions (**Materials and Methods, Supplementary Information**). A linear degradation rate of fluorescein was required to recapitulate the steady-state experimental data (**Fig. S1**).

### Investigating unidirectional bacterial signaling

Microbes communicate via chemical signals to monitor their population size, control gene expression and allocate resources within multispecies communites^30–33^. The spatial proximity of microbial populations is a critical variable shaping bacterial communication by dictating the local concentrations of quorum sensing molecules. We investigated the impact of spatial separation on the dynamics of quorum-sensing communication between bacterial populations. We constructed a synthetic community consisting of an *E. coli* sender strain that produces a quorum sensing signal (3-oxo-C6-HSL or AHL) that activates the expression of a fluorescent reporter (RFP) in an *E. coli* receiver strain (**Fig. 1b**). The sender strain harbored an arabinose-inducible AHL synthetase (LuxI) transcriptionally fused to a GFP fluorescent reporter and the receiver strain contained an aTc-inducible AHL receptor (LuxR) and a LuxR-regulated RFP fluorescent reporter **(Fig. S2)**.

We seeded MISTiC growth chambers with the sender and receiver strains and monitored their gene expression using time-lapse fluorescence microcopy and an automated image analysis pipeline (**Materials and Methods**). After an initial growth phase, arabinose was administered into the device to induce expression of LuxI and GFP in the sender strain (Experiment 1, **Table 1**). The interaction channel length did not significantly alter the response time of bacterial communication (**Fig. S3a**). The GFP steady-state did not vary significantly across interaction channel lengths, demonstrating the uniformity of arabinose concentration across the device (**Fig. 1c**). By contrast, the receiver’s steady-state RFP expression decreased as a function of distance from the sender growth chambers (**Fig. 1d**). Increasing the interaction channel length from 25 μm to 250 μm yielded a ~75% decrease in the receiver’s steady-state RFP expression levels. In sum, these results indicate that the spatial positioning of microbial populations can be encoded in the pattern of gene expression and MISTiC can be used to analyze the variation in gene expression across spatial separation.

### A dynamic model for inter-strain communication across distance

Building on our chemical diffusion model (**Supplementary Information**), we constructed a computational model of inter-strain bacterial communication across distance to understand the role of molecular factors on distance-dependent diffusion-mediated interactions in microbial communities. Mirroring the fluorescent dye model, diffusion across the interaction channels is represented as a one-dimensional process (**Materials and Methods, Supplementary Information**) (**Fig. 2a**). In the sender strain, the model captures the concentrations of arabinose (*ara*), GFP mRNA (*GFP*_*m*_), GFP protein (*GFP*_*p*_), LuxI mRNA (*LuxI*_*m*_), LuxI protein (*LuxI*_*p*_) and AHL (*AHL*). The species in the receiver strain include RFP mRNA (*RFP*_*m*_), RFP protein (*RFP*_*p*_), LuxR protein (*LuxR*_*tot*_) and the activated receptor (*LuxRAHL*) consisting of a complex of AHL bound to LuxR. The model includes time delays for arabinose transport and sequential assembly reactions for the proteins *GFP*_*p*_, *LuxI*_*p*_ and *RFP*_*p*_^34^. We used a genetic algorithm to estimate the parameters based on time-series fluorescent reporter measurements (**Materials and Methods**).

**Figure 2.**
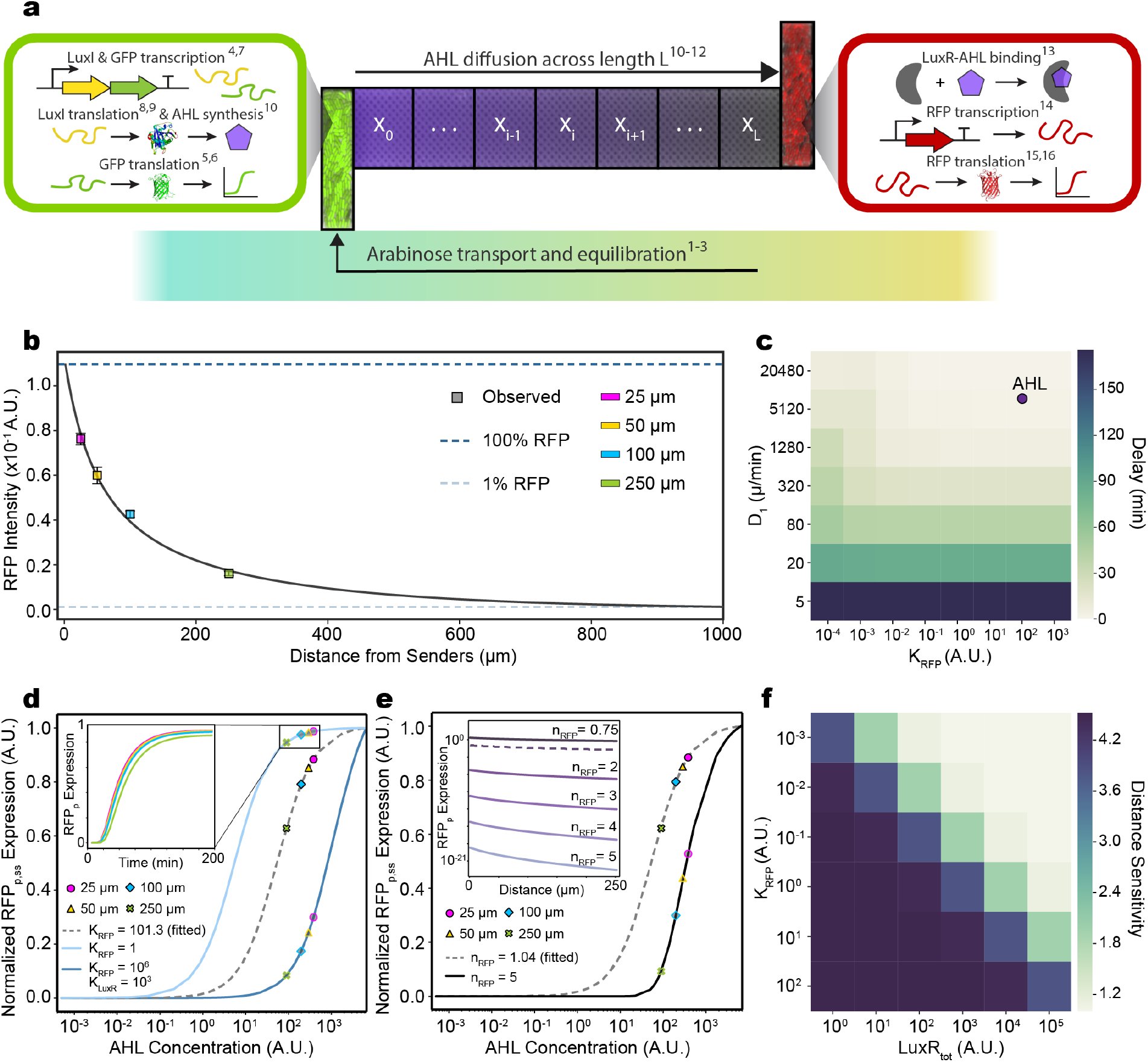
Computational model of inter-strain communication identifies key parameters influencing distance-dependent gene expression responses. **(a)** Model schematic depicting the physical and biological processes represented by the model equations (**Supplementary Information**). **(b)** The model *RFP*_*p*_ steady-states decrease as a function of distance from the sender population. The blue dashed line represents the maximum *RFP*_*p*_ steady-state concentration for a 2 μm interaction channel. The gray dashed line denotes 1% of the maximum steady-state *RFP*_*p*_ concentration. Data points (square) represent experimental measurements and error bars denote one standard deviation from the mean. **(c)** Heat map of the simulated *RFP*_*p*_ time delays for diffusible molecules spanning a broad range of diffusion rates (*D*_*1*_) and binding affinities of *LuxRAHL* to the *RFP*_*p*_ promoter (*K*_*RFP*_). The circle represents the estimated parameters based on experimental data. **(d)** *RFP*_*p*_ steady-state dose-response as a function of AHL for different *K_RFP_* values. The dashed curve represents the dose-response of the parameterized model based on the experimental data. Inset denotes representative *RFP*_*p*_ simulations for different interaction channel lengths. The *RFP*_*p*_ steady-states for each interaction channel length (colored marker styles) were computed for *ara*=10. **(e)** Steady-state *RFP*_*p*_ dose-response as a function of *AHL* for two different *n*_*RFP*_ values in the model. The dashed curve represents the dose-response for the parameterized model based on experimental data. The *RFP*_*p*_ steady-states for each interaction channel length (colored marker styles) were computed for *ara*=10. Inset: relationship between the separation distance and the steady-state *RFP*_*p*_ concentration for a range of *n*_*RFP*_ values. **(f)** Heat map of distance sensitivity across a broad range of total LuxR concentrations (*LuxR*_*tot*_) and binding affinities of the activated LuxR complex to the *RFP*_*p*_ promoter (*K*_*RFP*_). Distance sensitivity is defined as the ratio of steady-state *RFP*_*p*_ concentration for a 25 μm to 250 μm interaction channel. Source data are provided as a Source Data file.

The model parameterized to experimental data is able to accurately recapitulate the temporal changes in GFP and RFP at different distances (**Fig. 1c,d**). The model predicts that the steady-state receiver gene expression is highly sensitive to variation in spatial separation less than 100 μm **(Fig. 2b)** and forecasts that 150 bacterial cells can transmit information across 1000 μm based on a minimum threshold in *RFP*_*p*_ (defined as 1% of the maximum *RFP*_*p*_ expression at 2 μm separation). In the model, increasing the separation distance from 25 μm to 250 μm resulted in a 2.3 min delay in the RFP response time (**Fig. S3b)**. This was consistent with the absence of a detectable time delay due to the 7 min experimental measurement time resolution.

The diffusion rate of AHL from the growth chamber into the main channel (*D_2_*) and the degradation rate of AHL (*γ*_*AHL*_) influence the AHL concentration gradient established in the interaction channel. We sought to investigate the effects of these parameters on the distance-dependent gene expression pattern. Setting *γ*_*AHL*_ and/or *D*_*2*_ to zero significantly altered absolute *RFP*_*p*_ steady-state concentrations and their relative differences across distance, indicating that the stability of the chemical signal and the physical properties of the environment can dictate the response of a microbial community to spatial separation **(Fig. S4)**.

The response time of networks regulated by quorum-sensing can be modulated to optimize resource allocation to changeable environmental conditions^35,36^. Therefore, we explored how the *RFP*_*p*_ response time depends on two key parameters: the diffusion constant through the interaction channel, *D*_*1*_, and the binding affinity of *LuxRAHL* to the RFP promoter, *K*_*RFP*_. The delay in *RFP*_*p*_ increases with decreasing *D*_*1*_ and remains relatively constant as a function of *K*_*RFP*_. At intermediate values of *D*_*1*_, the delay is inversely related to *K*_*RFP*_ (**Fig. 2c**). The estimated parameters for the sender-receiver consortium map to a regime that display small time delays, indicating that a that a large change in *D*_*1*_ is required to yield a measurable time delay across a ten-fold change in distance.

We hypothesized that the distance sensitivity of information transmission in a microbial community can be controlled by biochemical parameters in the receiver strain. We analyzed our model to identify these parameters and found that the binding affinities of LuxR to AHL (*K*_*LuxR*_), *LuxRAHL* to the RFP promoter (*K*_*RFP*_), as well as the steepness of the steady-state response to AHL (*n*_*RFP*_) are critical in determining the steady-state gene expression response of the receiver to AHL concentration gradients. We mapped these parameters to changes in *RFP*_*p*_ steady-states across different distances from the sender strain. Changing *K*_*LuxR*_ and/or *K*_*RFP*_ to its target promoter (**Fig. 2d**), or *n*_*RFP*_ (**Fig. 2e**) relative to their estimated values shifts the system between the linear and saturated regimes of the dose response to AHL. In the linear regime, the *RFP*_*p*_ steady-state exhibits larger fold changes as a function of separation distance (greater distance-dependent sensitivity) compared to the saturated regimes (lower distance-dependent sensitivity). These results suggest that circuits could be programmed to realize different spatial patterns by modifying ultrasensitivity^37,38^, the affinity of transcription factors to promoters or a chemical inducer^39^, or the concentration of molecular factors in the circuit **(Fig. 2f)**.

### Spatiotemporal parameters impact fidelity of information transmission

Microbes in nature are constantly confronted with temporal variations in environmental stimuli. The spatial organization of a community can influence its response to these environmental perturbations^40^. A common strategy in engineering is using periodic input signals to characterize the dynamic properties of a system. We therefore applied a periodic input to the sender-receiver consortium to understand how distance affects information transfer.

To predict the behavior of this system, we simulated square wave oscillations in arabinose (*ara*) concentration with a period of two hours in the model (**Fig. 3a**). Our results show that the steady-state amplitude and mean *RFP*_*p*_ decreases as a function of increasing spatial separation. To test the model predictions, we performed MISTiC experiments using the sender-receiver consortium and alternated arabinose between 0% and 0.1% with a period of two hours (**Movie S1**) (Experiment 2, **Table 1**). The periods of GFP and RFP were synchronized with the arabinose input and therefore did not vary with distance (**Fig. S5a**). In response to the oscillatory signal, both GFP and RFP mean intensities increased over time and reached a steady-state oscillatory phase (**Fig. 3b,c**). The GFP mean and amplitude did not vary across distance (**Fig. 3b, Fig. S5b**). Mirroring the model prediction, the RFP mean, and amplitude decreased as a function of distance (**Fig. 3c, Fig. S5b**). At steady-state, the oscillatory activation and decay response times required were approximately equivalent for both GFP and RFP (30-40 min) (**Fig. S5c,d**). These response times are similar to the exponential phase doubling time of *E. coli* in LB media, suggesting that cell growth/division dictated the oscillatory timescale.

**Figure 3.**
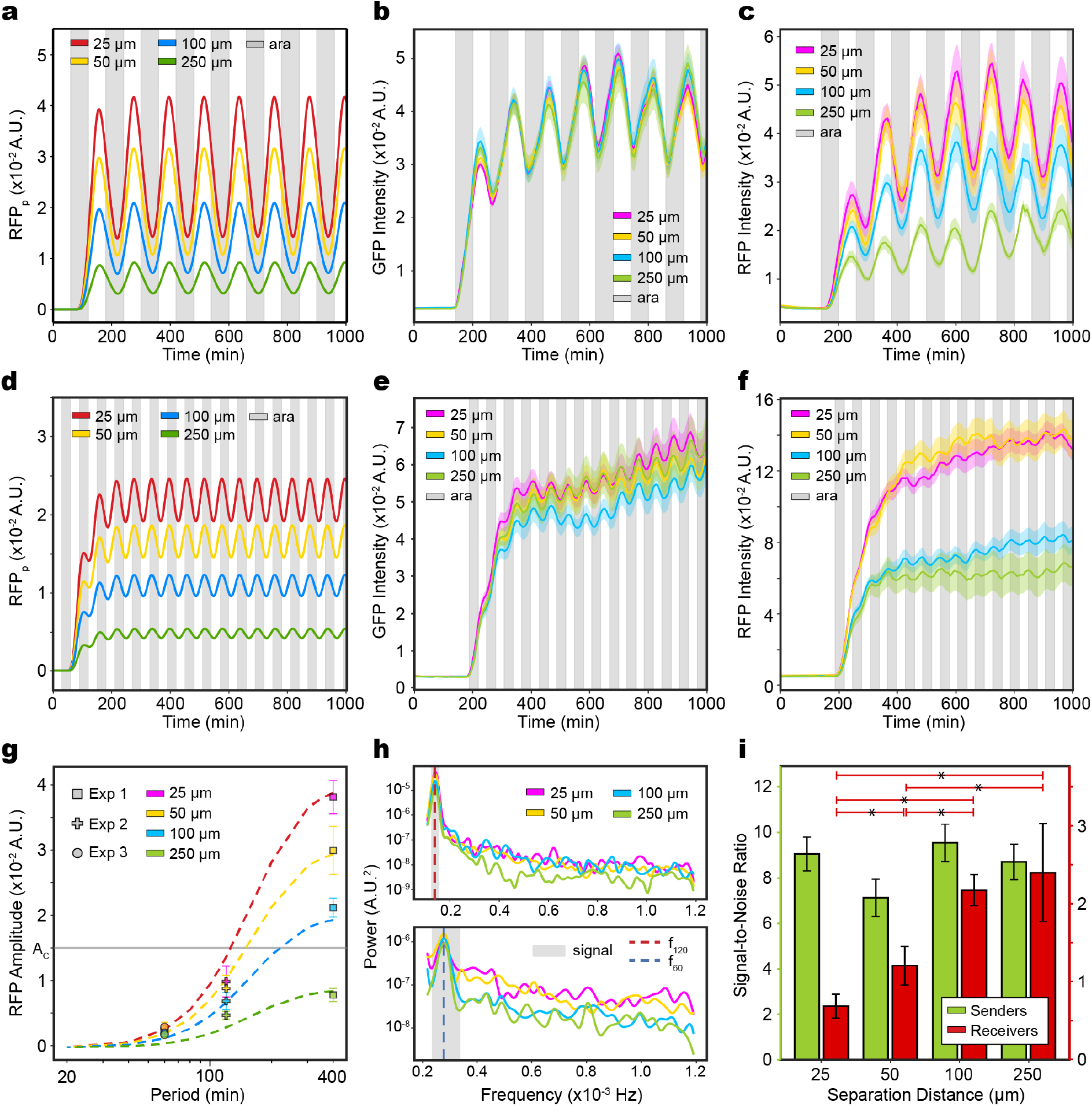
The frequency of a periodic input determines the effect of distance on information transmission in the sender-receiver consortium. **(a)** Model simulations of *RFP*_*p*_ expression across distance in response to a periodic arabinose input with a period of 2 hr. Gray shaded regions denote the presence of arabinose (**Table S3**). **(b)** GFP (sender strain) as a function of time in response to an oscillatory arabinose input with a period of 2 hr. Gray shaded regions denote the presence of 0.1% arabinose. Shaded regions represent one standard deviation from the mean. **(c)** RFP as a function of time in response in response to arabinose oscillations with a period of 2 hr. **(d)** Model simulations of *RFP*_*p*_ for different distances separating sender and receiver in response to square-wave oscillations in arabinose for a period of 1 hr. **(e)** GFP over time in response to an oscillatory arabinose input with a period of 1 hr. **(f)** RFP in response to an oscillating arabinose input with a period of 1 hr. **(g)** RFP amplitude as a function of the arabinose input period for different distances between the sender and receiver in the model (dashed lines). Square, plus, and circular data points represent the amplitudes in the step-response, 2 hr forced oscillator and 1 hr forced oscillator experiments, respectively. The gray horizontal line is a representative amplitude threshold (A_C_). **(h)** *Top:* RFP power spectra for a 2 hr arabinose period (red dashed line) for 395-1536 min. The gray region denotes a bandwidth of ±10 min around the expected input frequency (signal). *Bottom:* RFP power spectrum for a 1 hr arabinose period (blue dashed line) for 500-1249 min. The gray region denotes a bandwidth of ±10 min around the expected input frequency (signal). **(i)** Signal-to-noise (SNR) ratios computed using the power spectra for GFP or RFP at each interaction channel length (**Materials and Methods**) for the one-hour forced oscillator experiment. Error bars represent one standard deviation from the mean. Horizontal lines denote a statistically significant difference (P < 0.05) based on bootstrapped hypothesis testing. Source data are provided as a Source Data file.

We next explored whether information transfer across distance is corrupted by stochastic processes within the cell in response to a faster periodic input signals^41^. To investigate the temporal limitations of bacterial signal communication, we quantified the response of the sender-receiver consortium to a periodic arabinose input with a period of one hour. This period was predicted to push the limits of information transmission in the community based on the 30-40 min response time of the system (**Fig. S5c,d**). Simulations of square wave arabinose oscillations with a one-hour period yielded distance-dependent steady-state *RFP*_*p*_ responses with lower means and amplitudes compared to an input period of two hours (**Fig. 3d**). We characterized the response of the sender-receiver system in MISTiC to arabinose concentrations that alternated between 0% and 0.1% with a one-hour period (Experiment 3, **Table 1**). Sender GFP expression exhibited a steady-state oscillatory response and the GFP mean did not vary across interaction channel lengths (**Fig. 3e**). By contrast, RFP displayed temporal variability around the mean at steady-state as well as a distance-dependent change in the steady-state fluorescence intensity (**Fig. 3f**).

Amplitude fold change detection is an important property of sensory systems in diverse organisms^42^. A decreasing trend in the amplitude with increasing distance and higher input frequency was confirmed by simulations of *RFP*_*p*_ expression that varied the arabinose period over a wide range. The experimentally observed trend of decreasing amplitude as a function of input frequency mirrored the model predictions (**Fig. 3g**). Together, our results demonstrate that distance can establish a low-pass filter, where a minimum threshold in the *RFP*_*p*_ amplitude at a fixed period is detected at 25 μm but not 250 μm. Increasing the ultrasensitivity of RFP transcription (*n*_*RFP*_) augments the differences in RFP amplitude across increasing distance as a function of the input period (**Fig. S6a**), indicating that ultrasensitivity can enhance the switch-like response of the low-pass filter. In contrast to the two hour forced oscillator, the steady-state RFP amplitudes in response to a periodic input with a one hour period did not vary across distance (**Fig 3f,g**). This result suggests that a one-hour input period is approaching a critical frequency corresponding to the loss of information transmission in diffusion-mediated communication across distances of tens of microns^43^.

Communication between physically separated bacterial populations is impacted by extracellular noise due to diffusion^44^, as well as noise from intracellular processes such as transcriptional^45^ or translational bursting^46^. Such sources of noise can have a larger impact on gene expression dynamics in response to environmental fluctuations on faster timescales. We thus investigated how information in the periodic input signal is encoded in the frequency domains of the steady-state gene expression. The power spectrum represents how the variance in gene expression is distributed across frequencies. The power spectra for GFP and RFP displayed prominent peaks at the frequency of the input signal for both experiments (**Fig. 3h, Fig. S6b,c**). The RFP power spectrum peak at the input frequency decreased with distance in the two-hour forced oscillation experiment (**Fig. 3h**, top), reflecting the pattern in amplitudes across distance. In the one-hour oscillator experiment, the RFP power spectrum decreased with interaction channel length for frequencies larger than the signal bandwidth (**Fig. 3h**, bottom).

To evaluate the fidelity of information transmission across distance in these experiments, we defined the signal-to-noise ratio (SNR) as the total power of the input signal bandwidth divided by the total power across all frequencies greater than the signal bandwidth (**Materials and Methods**). The GFP SNR did not vary as a function of distance in either of the forced oscillator experiments (**Fig. 3i, Fig. S6d**). In the two-hour forced oscillator experiment, the RFP SNR is dominated by the power of the input signal and thus the SNR was inversely proportional to distance (**Fig. S6d**). Notably, the RFP SNR increased with distance in the one-hour forced oscillator experiment (**Fig. 3i**), indicating that the fidelity of information transmission was enhanced at longer distances.

In the regime of a critical input frequency^43^, the fidelity of inter-strain communication is diminished at short distances due to elevated noise whereas diffusible signals have a limited spatial range over long distances^19^. Therefore, our data suggests that the reliability of information transmission in quorum-sensing mediated communication varies non-monotonically with distance. Together the data shows that distance can function as a low-pass filter to allow cells to selectively respond to prolonged environmental fluctuations. Above a critical input frequency where cellular noise dominates, our data suggests that spatial separation can modulate a trade-off between the reliability of information transmission and the output expression level.

### Feedback loops impact oscillatory dynamics in response to spatial perturbations

The intracellular networks mediating microbial interactions can be complex and comprise positive and negative feedback loops and bidirectional communication. We sought to understand the effects of spatial separation on the gene expression dynamics in a microbial community wired together by bidirectional quorum-sensing communication and positive/negative feedback loops^47^. In this system, an *E. coli* activator strain produces C4-homoserine lactone (C4-HSL), which induces the enzymatic synthesis of 3-OHC14-HSL in an *E. coli* repressor strain (positive inter-strain interaction). The activator displays a positive feedback loop by self-regulating the circuit controlling C4-HSL production (**Fig. 4a**). The signal 3-OHC14-HSL produced by the repressor strain induces the expression of a quorum-quenching lactonase *aiiA* in the activator and repressor strains, which degrades both signals and thus inhibits circuit activity in the repressor (negative feedback loop) and activator (negative inter-strain interaction). An identical promoter driving the synthetases in the activator and repressor strains was used to drive the expression of CFP and YFP, respectively, to quantify circuit activity dynamics.

**Figure 4.**
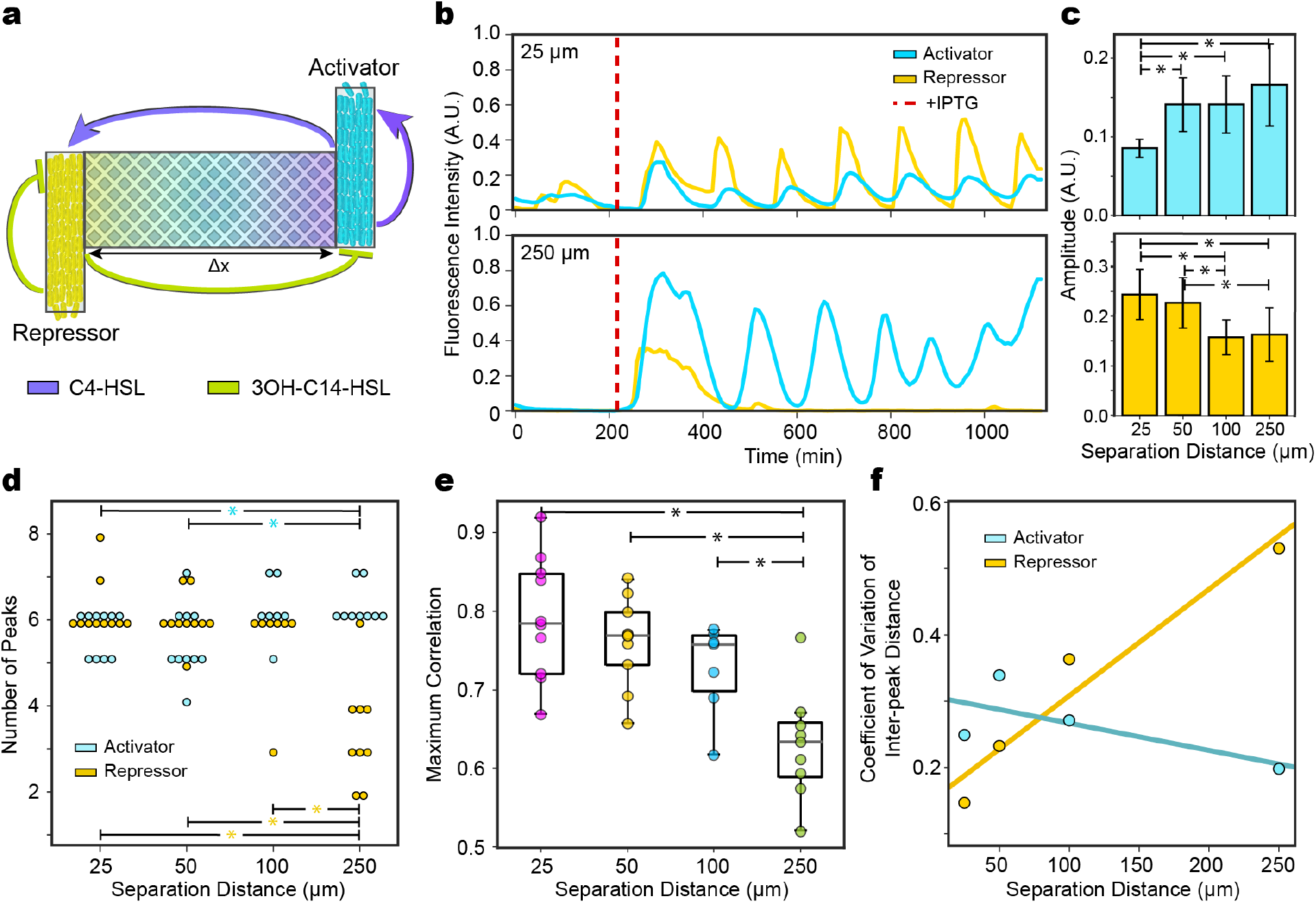
Inter-strain network topology and dual-feedback loops mediated by two orthogonal signals determine the effects of distance on a distributed oscillator consortium. **(a)** Network schematic of activator and repressor strains in the MISTiC device. The activator and repressor exhibit a bidirectional positive/negative interaction topology wherein the signaling molecules C4-HSL and 3OH-C14-HSL control circuit activities. The activator and repressor exhibit positive and negative feedback loops, respectively. **(b)** Representative normalized CFP and YFP fluorescence intensities as a function of time in the 25 μm (top) and 250 μm (bottom) distances. Red line indicates the time of induction with 1 mM IPTG. **(c)** Amplitude of peaks in the mean-subtracted CFP (top) and YFP (bottom) fluorescence intensities across distances. Error bars represent one standard deviation from the mean. Horizontal lines with stars denote a statistically significant difference (P < 0.05) based on an unpaired t-test. **(d)** The number of peaks as a function of distance for activator and repressor strains. Horizontal lines denote a statistically significant difference (P < 0.05) based on an unpaired t-test. **(e)** Maximum cross-correlation between paired CFP and YFP time-series measurements for each distance. Horizontal lines denote a statistically significant difference (P < 0.05) based on an unpaired t-test. **(f)** The coefficient of variation of the inter-peak distances (phase drift) as a function of separation distance for activator and repressor strains. Lines represent linear regression fits to the data. Source data are provided as a Source Data file.

The reporters CFP and YFP displayed oscillations for an interval of time across the majority of conditions in the MISTiC device (**Fig. 4b, Fig. S7, Movie S2**) (Experiment 4, **Table 1**). Paired growth chambers exhibited synchronized oscillations whereas unpaired growth chambers were not synchronized, indicating that diffusion of molecules through the interaction channels enhanced the coupling between the distributed oscillators. The CFP amplitude increased as a function of distance whereas the YFP amplitude displayed the reverse trend (**Fig. 4c**), signifying a reduced strength of the inter-strain interactions as a function of distance. The number of activator peaks detected in each growth chamber moderately increased, whereas the number of repressor peaks substantially decreased with separation distance (**Fig. 4d**). Therefore, the stability of oscillations in the repressor was highly sensitive to variations in the interaction range, whereas the oscillations in the activator displayed robustness to variations in this parameter. In specific 100 μm and 250 μm conditions, oscillations in the activator were maintained for a period of time after the repressor had stopped oscillating (**Fig. S7c,d**). The mean activator expression increased as a function of time for several replicates in 100 μm and 250 μm conditions, indicating that the activator would eventually approach a constitutive ON state as distance increased. Thus, our data suggests that the activator amplitudes vary non-monotonically with distance. The amplitudes are diminished by enhanced repression at short distances, increased at intermediate distances (**Fig. S7a,b**), and vanish at long distances as the activator expression approaches a constitutive ON state. Conversely, the repressor amplitudes decrease with distance and display an abrupt loss of oscillatory behavior at a critical distance threshold between 100-250 μm.

We next investigated the cross-correlation between the fluorescent reporters in the sender and receiver strain to quantify the effect of distance on the coupling of inter-strain gene expression dynamics. The maximum cross-correlation and time lag of the maximum cross-correlation decreased and increased with the length of the interaction channel, respectively (**Fig. 4e**, **Fig. S8a**), showing that distance diminished the coupling between the oscillators. The distribution of interpeak distances provides information about the variability in the oscillatory response and is an indicator of phase drift^48^. The coefficient of variation of the interpeak distance distribution increased by more than 3-fold for the repressor strain and was less variable across distance in the activator (**Fig. 4f, Fig. S8b,c**). In sum, the effects of distance on the oscillatory dynamics of the activator and repressor strains were notably different, indicating that the system’s feedback loops could be wired to enhance the robustness of oscillations across a range of distances. Previous studies have shown that negative enhances system stability, whereas positive feedback is associated with signal amplification, multi-stability and runaway behavior^49–51^. By contrast, positive and negative feedback exhibited the opposing roles for the distributed oscillator community by enhancing and reducing the temporal robustness and stability of the oscillations to spatial perturbations, respectively.

### Deciphering the spatial and temporal modes of metabolic interactions

In microbial communities, growth and metabolic activities are driven by metabolite dynamics including competition over limiting resources, metabolite secretion and cross-feeding^5,7,8,26,28^. Indeed, metabolite exchange is a major force shaping microbial communites^52^. To investigate how spatial separation influences metabolic interactions in microbial communities, we studied a synthetic *E. coli* consortium composed of a phenylalanine (*ΔpheA*) and methionine (*ΔmetA*) auxotroph. These auxotrophs were selected because bioinformatics analyses predict that phenylalanine (F) or methionine (M) auxotrophy are prevalent in microbial communities^53^. Further, M and F are two of the most energetically costly amino acids to synthesize in *E. coli*, potentially providing a potential selective advantage for cross-feeding in specific environmental contexts^53^. To characterize the population dynamics of the community in batch culture and evaluate longer-term stability, the strains were combined at three initial ratios (50% *ΔmetA*, 50% *ΔpheA*, 90% *ΔmetA*, 10% *ΔpheA*, and the reciprocal), in minimal media lacking M and F and then serially transferred three times to fresh media over a period of 5.5 days (**Materials and Methods**). Irrespective of the initial strain proportion, the co-culture converged to a *ΔpheA* dominated steady-state (**Fig. S9a**). For a sustained 24 hr passaging period, the community exhibited a decreasing trend in OD_600_ and eventually collapsed after the third passage (**Fig. S9b**, top). However, community growth was rescued by switching the final passage to a 48 hr incubation time, suggesting that a minimum cell density was required to maintain community growth and stability (**Fig. S9b**, bottom). These data show that the frequency of environmental shifts is a major determinant of community stability in batch culture and both strains can maintain growth and stable coexistence over multiple passages with *ΔpheA* dominating the community.

We next studied the growth responses of the strains within the spatially and temporally controlled environment of MISTiC. The population-level growth rate can be inferred by the rate of dilution of an inducible and highly-expressed stable fluorescent reporter due to cell growth^41,54^. We used this method to determine the maximum growth rates of the *E. coli* auxotroph community expressing GFP (*ΔmetA*) and RFP (*ΔpheA*) separated by defined distances in the MISTiC platform (**Fig. 5a**) (**Materials and Methods**). The fold change in the maximum growth rate of each strain across interaction channel lengths was used to quantify a distance-dependent interaction strength. In media supplemented with all amino acids, our results showed that *ΔmetA* and *ΔpheA* exhibited similar average doubling times of 80 and 86 min (Experiment 5, **Table 1**), respectively, across all interaction channel lengths (**Fig. 5b,c, S10a,b,c,d**), indicating a neutral interaction network (**Fig. 5h**).

**Figure 5.**
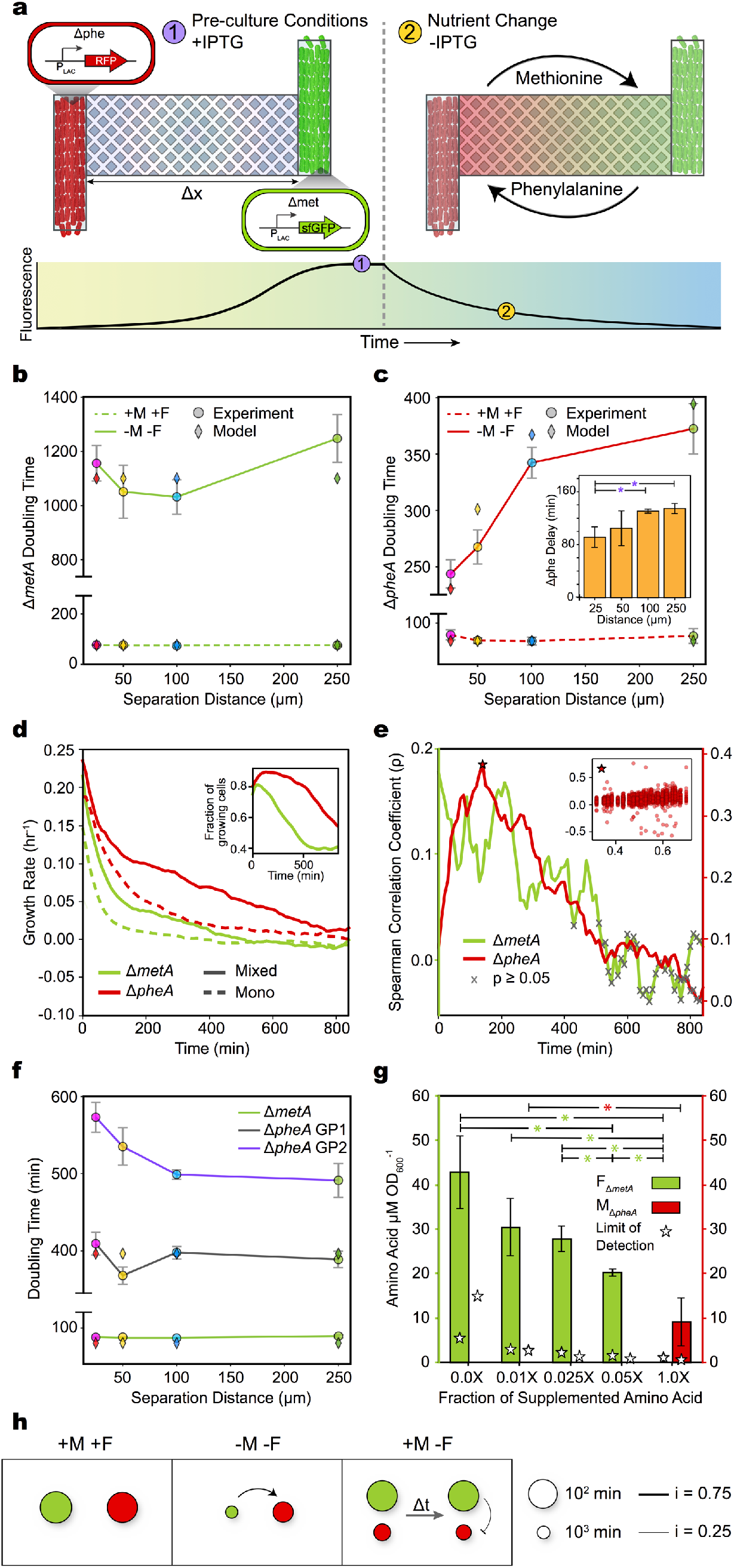
Spatial and temporal modes of amino acid cross-feeding in a synthetic *E. coli* consortium. **(a)** Schematic of the experimental design to infer the population-level growth rates of amino acid auxotrophs. The *E. coli* strains *ΔmetA* (IPTG-inducible GFP) and *ΔpheA* (IPTG-inducible RFP) are seeded into growth chambers in media supplemented with all amino acids and IPTG. At a specific time, the media is switched to remove IPTG and alter the amino acid concentrations. The rate of decay of the fluorescence intensity as a function of time can be used to infer the maximum growth rate in each growth chamber across distances. **(b)** Relationship between distance and the inferred population-level doubling times of *ΔmetA*. The solid and dashed lines represent absence and presence of methionine (M) and phenylalanine (F), respectively. Error bars represent one standard deviation from mean doubling times. Diamonds represent the model fit to the experimental data. **(c)** Relationship between distance and the inferred population-level doubling times for *ΔpheA*. The solid and dashed lines represent absence and presence of M and F, respectively. The horizontal lines denote a statistically significant difference (P < 0.05) based on an unpaired t-test. Diamonds represent the model fit to the experimental data. Inset: the time corresponding to maximum *ΔpheA* growth rate in media lacking M and F. **(d)** Relationship between time and the average growth rates of *ΔmetA* and *ΔpheA* in a mixed community (solid line) or monoculture (dashed line) in media lacking M and F. Inset: Fraction of cells with non-zero growth rates as a function of time. **(e)** Spearman correlation coefficient as a function of time between the fraction of the growth chamber occupied by the partner strain and the growth rate for individual cells (inset). The X symbol denotes correlations corresponding to P > 0.05. **(f)** Relationship between the distance and the minimum doubling time in the presence of M and absence of F. Growth phase 1 (GP1) and 2 (GP2) denote the two *ΔpheA* growth phases following the media switch. Diamonds indicate model fits to the *ΔmetA* and *ΔpheA* GP1 doubling times. **(g)** Concentration of M or F normalized by absorbance at 600 nm (OD_600_) in *ΔpheA* or *ΔmetA* filtered conditioned media. Stars indicate the limit of detection for each measurement. 1X amino acid fraction refers to 0.2 mM M and 0.4 mM F, respectively. **(h)** Inferred interaction networks based population-level minimum doubling times in physically separated MISTiC experiments. The size of the node is proportional to the maximum growth rate of each strain separated from the partner strain by 25 μm. Node diameters are scaled according to (10*ln(doubling time)^−1^)^2^. The edge widths are proportional to the distance-dependent change in growth rate D_d_ at 25 μm versus at 250 μm from the partner strain, where D_d_ = 1 − (D_25_D_250_^−1^). Source data are provided as a Source Data file.

We sought to determine the interaction network within a spatially separated, continuous flow environment in the absence of M and F (Experiment 6, **Table 1**) **(Fig. S11a,b)**. The *ΔmetA* strain exhibited a very slow doubling time (1128 min) that did not vary across distance (**Fig. 5b, Fig. S11a,c)**. In contrast, the doubling time for *ΔpheA* substantially increased with the separation distance, wherein a ten-fold increase in separation distance increased the strain’s doubling time by 54% (**Fig. 5c, S11b,d**). Therefore, the *ΔpheA* growth rate was highly sensitive to distance from the *ΔmetA* strain and not the reciprocal, highlighting a substantial difference in the strength of the interactions (**Fig. 5h**, center). Our data showed a 43 min time delay in the maximum growth rate of *ΔpheA* from the 25 μm to the 250 μm condition **(Fig. 5c**, inset), demonstrating that the transition from lag phase to growth was also distance-dependent.

We next investigated the growth of *ΔpheA* and *ΔmetA* in mixed conditions for comparison to the spatially separated context. A mixed culture containing equal proportions of *ΔmetA* and *ΔpheA* was introduced into the growth chambers in minimal media lacking M and F (Experiment 7, **Table 1**). Single-cell segmentation and tracking were performed to distinguish the strains within mixed communities and quantify the growth rates of single-cells (**Materials and Methods**) (**Fig. S12a**). The length of the interaction channels linking paired chambers did not contribute to the *ΔmetA* and *ΔpheA* growth rate variation (**Fig. S12b,c**). The average *ΔmetA* and *ΔpheA* growth rates within a chamber decreased over time but remained non-zero for the majority of the experiment (**Fig. 5d**). To evaluate the difference in each strain’s growth rate in the co-culture compared to monoculture, similar MISTiC experiments and analysis were performed for monocultures of *ΔmetA* (Experiment 8, **Table 1**) and *ΔpheA* (Experiment 9, **Table 1**) in minimal media lacking M and F. The co-culture growth rates were significantly higher than their respective monoculture conditions, demonstrating a mutual growth benefit in MISTiC. In addition, the percentage of growing cells across all growth chambers in the mixed condition was larger than 40% for both strains for the majority of the experiment (**Fig. 5d**, inset).

We computed the Spearman correlation between the growth rate of each strain and the fraction of the growth chamber occupied by the partner strain to quantify how the presence of the partner strain impacted growth rate (**Fig. 5e**, inset). Both strains exhibited statistically significant and non-zero Spearman correlations for the majority of the experiment, corroborating the presence of a mutualism. The *ΔpheA* strain exhibited a higher maximum Spearman correlation than the *ΔmetA* strain, corroborating a stronger dependence of *ΔpheA* on the abundance of *ΔmetA* than the reciprocal. Even though *ΔpheA* exhibited a higher growth rate than *ΔmetA*, the average ratio of the two strains reached a stable value (**Fig. S12d**). Since the percentage of dividing cells (**Fig. S12e**) was significantly lower than the percentage of growing cells (**Fig. 5d**, inset) for both strains, cell elongation was the dominant mode of growth in these conditions. Additionally, growth rates did not vary as a function of the position of single cells within the growth chambers (**Fig. S12f**). These findings illustrate the ability of MISTiC to resolve sub-population heterogeneities in growth based on single-cell data.

For the spatially separated community, we next examined whether the interaction network could be toggled by rescuing *ΔmetA* in minimal media supplemented with all amino acids except F (Experiment 10, **Table 1**). The doubling time of *ΔmetA* was similar to its doubling time in the +M, +F control experiment (Experiment 5, **Table 1**) and did not vary across distance (**Fig. 5f, S13a,d**). The rate of change of RFP fluorescence was biphasic, indicating that *ΔpheA* had two growth modes in this condition (**Materials and Methods, Fig. S12b,c**). The *ΔpheA* doubling times in the first growth phase did not change across distance, yielding a neutral interaction (**Fig. 5h**, **Fig. S13e**). The second *ΔpheA* growth phase displayed a moderately competitive growth trend across a ten-fold change in distance (**Fig. 5f,h, Fig. S13f**). The distance-dependent benefit from *ΔmetA* to *ΔpheA* was not detected in the presence of M, suggesting that the positive interaction is abolished by rapid growth of *ΔmetA*.

To probe the inverse relationship between the growth rate of *ΔmetA* and the magnitude of the benefit to *ΔpheA*, we measured the M and F concentrations in the producer strain’s supernatant across different concentrations of the rescuing amino acids (**Materials and Methods**). The F concentration in the *ΔmetA* supernatant was inversely proportional to the supplemented M concentration, consistent our MISTiC results showing that the greatest growth benefit to *ΔpheA* occurred when *ΔmetA* was slowly growing and metabolically active (**Fig. 5g**). Interestingly, the reverse trend was observed for *ΔpheA*, wherein M was detected only in the condition with the highest concentration of supplemented F. In all conditions, *ΔpheA* exhibited a higher growth rate, which could be explained by the high release rate of F by *ΔmetA* at low growth rates.

Our results demonstrated that *ΔmetA* and *ΔpheA* auxotroph strains have differential and context-dependent release rates of M and F. To integrate our findings into a quantitative framework, we developed a dynamic computational model to represent M and F biosynthesis, uptake, diffusion, and amino acid dependent growth rates of *ΔmetA* and *ΔpheA* (**Supplementary Information**). Mirroring the structure of the quorum-sensing model, M and F diffuse from the producer strain’s growth chamber across the interaction channel and into the recipient strain’s growth chamber. Consistent with our data, we assume that (1) growth is limited by the concentration of the amino acid that the auxotroph is deficient in producing, (2) release of F by *ΔmetA* is inversely proportional to its growth rate, (3) release of M by *ΔpheA* is proportional to its growth rate, and (4) the basal growth rate *ΔpheA* is larger than *ΔmetA* attributed to differences in the metabolic consequences of each mutation (**Fig. 5d**). The model was fit to the population-level growth rates in the physically separated experiments using a genetic algorithm (**Materials and Methods**) and was able to recapitulate the trends across a range of conditions (**Fig. 5b,c,f**), demonstrating that the model’s core assumptions were congruous with the data.

Finally, we tested how different concentrations of all amino acids impacted the interaction strengths within MISTiC (Experiments 11-13, **Table 1**). The *ΔpheA* strain displayed a decreasing trend in the distance-dependent interaction strength, defined as the ratio of its doubling times in the 25 μm to 250 μm condition, as a function of amino acid availability (**Fig. S14b,d,f,h**). An interaction strength trend across amino acid concentrations was not evident for *ΔmetA* (**Fig. S14,a,c,e,g**). Low amino acid concentrations eliminated distance-dependent growth rate response of *ΔpheA*, indicating that the growth benefit provided by *ΔmetA* was dependent on trace concentrations of F. In sum, our results suggest that the stability of an amino-acid auxotroph consortium is shaped by key variables including amino acid availability, population size, spatial arrangement of populations, temporal frequency of environmental fluctuations and environmentally regulated amino acid release rates. The change in growth as a function of interaction channel length indicates whether metabolites mediating microbial interactions are in a saturated (below a threshold to impact growth or not limiting for growth) or linear regime (limiting to growth). These insights cannot be obtained in a well-mixed batch culture and the operating regime of a microbial interaction could impact community stability and functions. In addition, MISTiC enabled the characterization of the dynamic features of the growth rate responses including biphasic growth (**Fig. S13b,c**) and delays in the timing of maximum growth (**Fig. 5c**, inset).

## DISCUSSION

We developed a microfluidic platform, MISTiC, to interrogate the effects of spatiotemporal parameters in microbial consortia. In our study, the distance between strains impacts the concentration of diffusible molecules, which in turn dictates the biological response including growth rate, metabolic activity or cell state decision-making. Therefore, changing the interaction channel length enables a quantitative mapping of the dose-response of growth rate or gene expression to the net environmental impact (produced or utilized diffusible mediators) of a partner strain. Using MISTiC, we can elucidate if a diffusion-mediated interaction is operating in a linear or saturated regime based on the change in growth or gene expression as a function of distance from the partner strain, which has implications for community stability and functions. Interactions operating in a saturated regime may promote community stability by enhancing robustness to environmental perturbations, whereas interactions operating in a linear regime may have the opposite effect of reducing stability and resilience to environmental shifts.

Key parameters including the binding affinity of transcription factors to their target promoter or chemical inducer and the ultrasensitivity of a promoter can shift the system between regimes that are sensitive (linear) or robust (saturated) to spatial variations. These biomolecular parameters can be tuned to spatially program division of labor (e.g. ecological niches or metabolic functions) for biotechnological applications. For instance, ultrasensitivity could be modulated to create defined spatial boundaries between distinct cell states^38^. In the sender-receiver quorum-sensing system, distance-dependent gene expression levels were produced by the AHL gradients established within interaction channels. Similarly, morphogen gradients in multicellular organisms dictate intracellular signaling and cellular differentiation^55^. Our results show that these chemical gradients are tunable by the spatial arrangement of populations and could be exploited to program complex pattern formation using microfabrication techniques and tools from synthetic biology^56^.

In natural environments, microbial communities are confronted with uncertain and changeable environmental conditions. We show that distance between community members establishes a low-pass filter that allows cells to selectively respond to signals that fluctuate on longer timescales. In addition, our data demonstrates that distance can improve the reliability of information transmission in response to specific input frequency ranges. Consistent with this result, theoretical work has shown that spatial averaging by diffusion can improve the precision of gene expression by reducing noise stemming from transcriptional bursting^57^. In addition, stochastic modeling predicts that diffusion of a quorum-sensing chemical signal could reduce gene expression noise^44^. Our results suggest that spatial positioning can tune the trade-off between the fidelity of information transmission and gain of a signal and could be exploited as a design feature in microbial community engineering. While increasing the expression level of circuit components can reduce stochastic fluctuations, it can also generate a metabolic burden to the cell^58^. In contrast, modulating the distance between populations is an alternative strategy to enhance the signal-to-noise ratio without incurring additional energetic costs.

Changing the degree of spatial separation between the activator and repressor strains in the dual-feedback oscillatory consortium yielded different outcomes on the oscillatory behaviors of each strain. For instance, the stable oscillatory behavior of the repressor was abruptly lost in the 250 μm condition, whereas oscillatory behavior persisted in the activator strain over a period of time. The notable difference in the robustness of the circuits to spatial perturbations highlights the critical role of circuit topology and feedback loops in determining system dynamics in spatially organized communities. Thus, molecular circuits can be wired together in different ways to amplify or reduce dynamic signals as a function of distance^19^. The activator and repressor display self-activation and self-repression, respectively, suggesting that positive feedback enhances the robustness of information transmission to changes in spatial separation, whereas negative feedback has the reverse effect of enhancing the fragility of the system. The lag required to maximize the cross-correlation between the activator and repressor oscillatory responses increased with interaction channel length (**Fig. S8a**), whereas the sender-receiver consortium in response to the step-response of arabinose did not exhibit a time delay across distance (**Fig. S3a**). In this case, dual-feedbacks and bidirectionality of the interaction could lead to back-and-forth propagation of quorum-sensing signals, thus augmenting the time-delay between the oscillations.

The disparity in *ΔpheA* and *ΔmetA* growth rates in the absence of F and M could be explained by their differential amino acid release profiles as a function of the producer strain’s growth rate (**Fig. 5g**). In *E. coli*, F is either used for either protein synthesis or transported between the periplasm and cytosol^59^. However, M can be transformed into S-adenosyl-L-methionine, which acts as a central hub methyl donor intersecting many pathways^59^. This suggests that M is a critical resource for the cell and may be transformed into other molecules at a faster rate than F, resulting in differential amino acid release profiles. Auxotrophic cross-feeding has been proposed as a strategy to enhance coexistence and stability among members of a consortium^53,60,61^. Our results suggest that strain co-existence and community stability are highly dependent on a critical population size, amino acid availability, temporal perturbations and spatial context^1,2,5^. Therefore, stable coexistence may be difficult to achieve in real-world environments which are spatially heterogenous and temporally changeable.

MISTiC enables quantification of the effect of micron-level spatial separation on diffusion-mediated microbial interactions, in the absence of convection, transport or cell-to-cell physical contact, by monitoring growth, gene expression, cell size, morphology or emergence of cellular states as readouts. Coexistence of members of a microbial community can be difficult to achieve in batch or continuous culture due to variation in growth rates and competition for limited nutrients. The physically separated and connected growth chambers in MISTiC can maintain strain coexistence over long periods of time to study microbial interactions. Whereas metabolite exchange between populations cannot be easily observed within MISTiC, biosensors for specific diffusible compounds could be used as real-time indicators of microenvironments^62^. The media flow rate is a key parameter that can be manipulated to study how temporal fluctuations shape microbial interactions or recapitulate the flow rates of natural environments. To determine if interactions stem from physical contact or diffusible compounds, a mixed chamber could be added to the device for comparison to spatially separated arrangements.

There are limited techniques to investigate small bacterial populations (~10^2^ cells), which exist in natural environments and can play important roles in human disease^63^. However, the size of the MISTiC growth chamber limits each strain’s impact on its environment, and thus the strength of interactions. To interrogate the contribution of population size to microbial interactions and community stability, future device designs will include growth chambers that vary in size. To investigate more complex communities and higher-order interactions, modified versions could be constructed to study three or more interacting populations. In sum, this experimental platform could be adapted to study a diverse repertoire of organisms and mechanisms of diffusion-mediated interactions over multiple length-scales and increasingly complex spatial landscapes. A detailed understanding of how defined spatial arrangements influence community functions and interaction networks will advance our understanding of the spatial organization of microbiomes inhabiting diverse natural environments.

## MATERIALS AND METHODS

### Microfluidic device fabrication

A three-layer device was designed in AutoCad that consisted of interaction channels, growth chambers, and main channels. The microfluidic master was pattered in three stages of photolithography using a micropattern generator (Heidelberg Instruments μPG 101). For the first layer, the silicon wafer was baked for 10 min at 200°C and spin coated at 4000 rpm using SU-8 2000.5 (MicroChem) to generate 0.5 μm height. This layer was exposed to the interaction channels at 58 mW with a 47% dwell time using a 4 mm writehead, followed by a post-exposure bake for 30 min at 95 °C. The second layer was spin coated at 3000 rpm using 26:1 mixture of SU-8 2000.5 to SU-8 3005 to produce 1.5 μm height. After aligning to the first layer, the wafer was exposed to the second patterning layer (growth chambers). Following an additional post-exposure bake, a third layer of SU-8 3025 photoresist was spin coated at 3000 rpm to generate 25 μm height. The wafer was exposed to the final layer consisting of the main channels, resistors, and inlets. Following a final post-exposure bake, the features were developed using SU-8 developer (MicroChem). The master was treated overnight with vapor phase (tridecafluoro-1,1,2,2-tetrahydrooctyl) trichlorosilane (Gelest) at room temperature. To fabricate each device, 7:1 mixture of polydimethylsiloxane (PDMS, Sylgard 184) to curing agent (Sylgard 184) was used to coat the master. After curing overnight at 100°C, the inlet and outlet holes were punched using a biopsy corer (WellTech). The surfaces were exposed to air plasma (Harrick Plasma PCD-32G) for 23 sec to ionize the surface of the device to bond to the glass coverslips (ThermoFisher). Finally, the surfaces were bonded and baked for 1 hr at 100 °C to seal the device channels. For each experiment, the microfluidic device was flushed with 0.5% Tween 20 (Sigma-Aldrich) to prevent cells from adhering to the device. To load the cells into the device, a vacuum pressure of 330 mmHg was applied load cells into the growth chambers.

### Dye gradient experiment

The chemical gradients in the interaction channels were analyzed by administering 10 μM fluorescein (Sigma-Aldrich) and water at a flow rate of 200 μL hr^−1^ into individual main channels. Paired growth chambers (n = 3) connected by each interaction channel length were continuously imaged using a 600 msec exposure time. Fluorescence and phase contrast Images were collected using a Ti-E Eclipse inverted microscope (Nikon) using the GFP filter (Chroma) 470nm/40nm (ex), 525/50nm (em). To analyze the images, the fluorescence in each growth chamber as well as 1 μm increments along the length of each interaction channel at steady-state were determined.

### Sender-receiver quorum-sensing experiments

Sender and receiver plasmids (**Fig. S2**) were constructed using standard Gibson assembly protocols using primers synthesized by Integrated DNA Technologies and verified by Sanger Sequencing (Functional Biosciences). The sender (A6c_LuxI_GFP^64^) and receiver (E2c_LuxR_RFP or pJH9-35) plasmids^65^ were transformed into *E. coli* strains BW27783^66^ and MG1655Z1^67^ (**Table 2**), respectively. An initial set of cultures were inoculated into LB media (Lennox, Sigma-Aldrich) containing 25 μg/mL chloramphenicol (Sigma-Aldrich) and cultured overnight at 37°C with shaking. After approximately 16 hr, 1 μL of the cultures were diluted into 3 mL LB media containing 25 μg/mL chloramphenicol and incubated at 37°C with shaking to early stationary phase (OD_600_ 0.7-1.1). Next, we measured the OD_600_ of these cultures and centrifuged 1 mL at 3500 x g. The supernatant was removed, and the pellet was resuspended to a final OD_600_ of approximately 20. Cells were loaded into the device such that each growth chamber had 2-3 cells at the beginning of the experiment. In each experiment, the device was connected to three syringes (5 mL) containing LB media supplemented with 25 μg/mL chloramphenicol, 0.1% Tween 20 (Sigma-Aldrich) and 62.5 ng/mL anhydrotetracycline hydrochloride (Cayman Chemicals) as well as a fourth syringe (5 mL) containing the same media supplemented with 0.1% arabinose (Sigma-Aldrich). During the microscopy experiment, the microfluidic device was incubated at 37°C in a custom designed temperature incubation chamber. The main channels were flushed at a rate of 300 μL hr^−1^ to wash away excess cells from the growth chamber. The flow rate of the inlet containing arabinose (I_22_, **Fig. 1**) and the corresponding inlet on the opposite side (I_11_) were set to 10 μL hr^−1^ to prevent cell growth and clogging within the inlet and resistor and reduce pressure differences across the device. Fluorescence and phase contrast images were collected using a Ti-E Eclipse inverted microscope (Nikon) every 7 min at 21 different positions. Fluorescence was imaged using the following filters (Chroma): GFP: 470nm/40nm (ex), 525/50nm (em) or RFP: 560nm/40nm (ex), 630/70nm (em). The device was incubated for a period of time to allow cells to grow and divide. After the growth chambers had filled with cells, the media was switched to test conditions described in **Table 1**. For Experiment 1 (**Table 1**), the arabinose inlet (I_22_, **Fig. 1**) and the corresponding inlet on the opposite side (I_11_, **Fig. 1**) were switched to 200 μL hr^−1^ and the flow rate through the remaining inlets (I_12_, I_21_) were set to 0 μL hr^−1^. The forced oscillation experiments (Experiment 2-3, **Table 1**) used 10 mL syringes to extend the length of the experiment. Flow rates of the 0.1% arabinose (inlet I_22_) and 0% arabinose (inlet I_21_) were alternated out of phase between 200 μL hr^−1^ and 0 μL hr^−1^ for a period of time. One of the receiver inlets (I_11_) flowed continuously at a rate of 200 μL hr^−1^ and the other inlet (I_12_) was set to 0 μL hr^−1^ for the duration of the experiment.

**Table 2.** Strains used in study. The * symbol indicates plasmids that were constructed for this study. All other constructs were derived from the indicated references.

| Strain identifier | Strain background | Plasmid(s) | Reference |
| --- | --- | --- | --- |
| Sender | BW27783 | A6c_LuxI_GFP* | 66 |
| Receiver | MG1655z1 | E2c_LuxR_RFP* | 67 |
| Activator | CY027 (E. coli ΔlacI<br>ΔaraC ΔsdiA Ptrc*-<br>cinR Ptrc*-rhIR)<br>Addgene #72402 | pC220 (Addgene<br>#65877) and pC239<br>(Addgene #65953) | 47 |
| Repressor | CY027 (E. coli ΔlacI<br>ΔaraC ΔsdiA Ptrc*-<br>cinR Ptrc*-rhIR)<br>Addgene #72402 | pC236 (Addgene<br>#65951) and pC239<br>(Addgene #65953) | 47 |
| ΔmetA | BW25113 | A6c_GFP* | 68 |
| ΔpheA | BW25113 | A6c_RFP* | 68 |

### Dual-feedback oscillator experiments

The *E. coli* strain CY027 was transformed separately with plasmids pC220 and pC239 or pC236 and pC239 to construct the activator and repressor^47^, respectively using a standard chemical transformation protocol (**Table 2**). Overnight cultures were inoculated into LB media (Lennox) containing 50 μg/mL kanamycin and 100 μg/mL spectinomycin and incubated overnight at 37°C with shaking. After approximately 16 hr, 1 μL of the overnight cultures were diluted into 3 mL LB media and incubated at 37°C with shaking to early stationary phase (OD_600_ 0.7-1.1).

Cells were loaded into the device following the procedure specified above. Following cell loading, the microfluidic chip was placed in the custom designed temperature incubation chamber at 37°C. All four inlets were connected to syringes (10 mL) containing LB media with kanamycin (50 μg/mL), spectinomycin (100 μg/mL) and 0.1% Tween 20. Syringes connected to inlets I_22_ and I_11_ also contained 1 mM isopropyl ß-D-1-thiogalactopyranoside (IPTG) (Sigma).

The cells were initially grown in the device at 37°C with inlets I_12_ and I_21_ flowing at 200 μL hr^−1^, and inlets I_22_ and I_11_ flowing at 10 μL hr^−1^ to prevent cell growth and clogging. Phase contrast and fluorescence images were collected every 7 minutes at 21 different positions. Once the growth chambers were filled with cells (**Table 1**), the inlets (I_12_ and I_21_) containing the pre-culture media were set to 0 μL hr^−1^ the inlets (I_11_ and I_22_) containing the test media were set to 200 μL hr^−1^.

### Amino acid auxotroph experiments

*E. coli* strains *ΔmetA*^68^ and *ΔpheA*^68^ (**Table 2**) were transformed with plasmids A6c_GFP^69^ and A6c_RFP^69^, respectively using a standard chemical transformation protocol. The plasmids harbored an IPTG inducible fluorescent reporter. An initial set of cultures were inoculated into LB media (Lennox) containing chloramphenicol (25 μg/mL) and incubated overnight at 37°C with shaking. After approximately 16 hr, 1 μL of the overnight cultures were diluted into 3 mL of LB containing 25 μg/mL chloramphenicol and 1 mM IPTG (Sigma-Aldrich) and incubated at 37°C with shaking until early stationary phase (OD_600_ 0.7-1.1).

The cells were loaded into the device following the procedure outlined above. Following cell loading, the microfluidic chip was placed in the custom designed temperature incubation chamber at 37°C. The media always contained 1X MOPS Buffer (Teknova), 1X ACGU mix (Teknova), chloramphenicol, 0.1% Tween 20, 1.32 mM potassium phosphate dibasic (Teknova), and 0.2% glucose (Teknova), whereas the amino acid composition varied across experiments (**Table 1**). The amino acid solutions consisted of either EZ Amino Acids (AA, Teknova) or a modified amino acid solution (AA*) (**Table 1**). The AA* solution consisted of 0.4 mM L-asparagine (VWR), 0.01 mM calcium pantothenate (VWR), 0.2 mM L-histidine (VWR), 10 mM L-serine (VWR), 0.8 mM L-alanine (Fisher Scientific), 0.4 mM L-lysine (Fisher Scientific), 0.1 mM L-tryptophan (Fisher Scientific), 0.4 mM L-aspartic acid (Dot Scientific), 0.1 mM L-cysteine (Dot Scientific), 0.8 mM L-glycine (Dot Scientific), 0.4 mM L-isoleucine (Dot Scientific), 0.8 mM L-leucine (Dot Scientific), 0.01 mM para-amino benzoic acid (Dot Scientific), 0.4 mM L-proline (Dot Scientific), 0.4 mM L-threonine (Dot Scientific), 0.6 mM L-valine (Dot Scientific), 5.2 mM L-arginine (Sigma), 0.01 mM di-hydroxy benzoic acid (Sigma), 0.6 mM L-glutamic acid (Sigma), 0.01 .mM para-hydroxy benzoic acid (Sigma), 0.01 mM thiamine (Sigma), 0.2 mM L-tyrosine (Sigma) and 0.6 mM L-glutamine (Acros Organics). In minimal media supplemented with AA*, varying concentrations of methionine (Dot Scientific) and/or phenylalanine (Dot Scientific) were added. The AA amino acid solution consisted of all components in AA* plus 0.2 mM methionine and 0.4 mM phenylalanine.

The cells were grown for a period of time at 37°C with 1 mM IPTG prior to the media switch as described in **Table 1** to fill the growth chambers. Phase contrast and fluorescence images were collected every 10 minutes at 21 different positions. After the growth chambers were filled with cells, the inlets (I_12_ and I_21_) containing the pre-culture media were set to 0 μL hr^−1^ and the inlets (I_11_ and I_22_) containing the test media were set to 200 μL hr^−1^.

### Amino-acid measurements

The *ΔmetA* and *ΔpheA* strains were inoculated into LB (Lennox) containing chloramphenicol (25 μg/mL) and grown overnight at 37°C with shaking. After 16 hours, 10 μL of the cultures were transferred into 3 mL of fresh LB containing chloramphenicol (25 μg/mL) and incubated at 37°C with shaking until early stationary phase (OD_600_ 0.7-1.1). Immediately following, the cultures were centrifuged at 3500 x g for 5 min, supernatant was removed and the cells were inoculated into MOPS EZ Rich Defined Medium lacking M and F at an initial OD_600_ of 0.05. For the *ΔmetA* strain, 0, 2, 5, 10 or 200 μM M was added to the media. For *ΔpheA* strain, 0, 4, 10, 20 or 400 μM F was added to the media. The cultures were incubated at 37°C with shaking for 3 hr. After recording the OD_600_ of each culture, the cells were centrifuged at 3500 x g for 10 minutes, the supernatant was filtered by a 0.2 μm filter (GE Healthcare) and the concentrations of M or F were measured with a fluorometric assay kit (BioVision) or by liquid chromatography-mass spectrometry (LC-MS) respectively. Concentrations of M in the filtered conditioned media of *ΔpheA* cultures were measured with a fluorometric methionine assay kit (BioVision) with a 0.5 μM limit of detection. Raw fluorescence measurements were converted to methionine concentrations using a standard curve.

The analysis of F concentrations in the filtered conditioned media of *ΔmetA* was performed on a Shimadzu LC-MS2020. All solvents and reagents used the analysis were HPLC grade or higher quality. Methanol and formic acid used for the measurements was sourced from Fisher Scientific and Acros Organics, respectively. Water was prepared in house with a Millipore Milli-Q water purification system. Separations were performed at 40°C on a Discovery BIO wide pore C5-5 column (15 cm × 2.1 mm × 5 μm) from Millipore-Sigma with a paired Supelguard (2 cm × 4 mm × 5 μm) guard column. The running buffer was a binary gradient of water with 0.1% v/v formic acid (Buffer A) and methanol (Buffer B) according to the following protocol: 4 minutes at 5% B, a linear gradient from 5% to 20% for 4 minutes, a linear gradient from 20% B to 95% B for 2 minutes, 2 minutes at 95% percent B, a linear gradient from 95% B to 5 % B for 2 minutes, equilibrating the at 5% B for 6 minutes. The total flow rate was 0.2 ml min^−1^. Under these conditions, methionine and phenylalanine eluted at 3.8 minutes and 5.8 minutes, respectively. The ion source was operated in ESI mode with a cone voltage of 4.5 kV, the interface was held at 400°C and the desolvation line at 250°C. The dry nitrogen was supplied to the nebulizer at 1.5 L min^−1^ and drying gas at 15 L min^−1^. The mass spectrometer was run in selective ion monitoring (SIM) for monitoring m/z 150 for methionine and m/z 166 for phenylalanine with a scan time of 1 second. Standards were prepared for each run by adding known concentrations of methionine and phenylalanine to fresh media. The standard curve was run before and after the sample batch and each sample was run twice for technical replicates.

### Auxotroph community batch culture experiment

Separate culture tubes containing LB with chloramphenicol (25 μg/mL) were inoculated with *ΔmetA* or *ΔpheA* and incubated overnight at 37°C with shaking. After 16 hours, the cultures were diluted into 5 mL of EZ Rich Medium (Teknova) containing chloramphenicol (25 μg/mL) and 1 mM IPTG and lacking M and F at a final OD_600_ of 0.05. The ratio of *ΔmetA* to *ΔpheA* was 10:1, 1:1, or 1:10 (n = 3, for each starting ratio). The cultures were incubated at 37°C with shaking for at least 24 hours before transferring the community to fresh media using a 1:100 dilution. At this transfer time, the OD_600_ of each culture was measured and a 2 μL sample was spotted onto a glass slide for cell counting with microscopy (20X magnification) on a Nikon Eclipse Ti. Four images comprising four distinct fields of view were taken of each sample and each image was a composition of phase contrast, GFP and RFP channels. Subsequently, ImageJ was used to extract the number of *ΔmetA* cells from the GFP channel and *ΔpheA* cells from the RFP channel for each image.

### Population-level image analysis

For Experiments 1,2,5,6,10-13 (**Table 1**), individual growth chambers were segmented in DeepCell^70^. Five neural networks were trained on 21 randomly selected images as well as binary masks (made using FIJI image analysis software^71^) that specified the growth chamber positions. The trained model was used to analyze the remaining microscopy images. The results of the trained networks (2-5 depending on segmentation accuracy) were averaged to improve segmentation accuracy.

For Experiments 3 and 4 (**Table 1**), growth chambers were segmented using custom code in Python that aligned each growth chamber across all time points. A binary mask denoting the growth chambers was applied to all time points. There was a negligible difference in the fluorescence time-series for the DeepCell and alignment methods.

In each analyzed image, custom code (Python) was used to label the binary mask with the growth chamber positions and total areas and compute the average fluorescence intensity of each growth chamber. Segmented regions less than or greater than 1000 and 3500 pixel area were eliminated from the data set. Specific criteria were used to eliminate outliers from the data sets including (1) infrequent pressure fluctuations leading to loss of cells from the growth chambers, (2) device bonding issues leading to collapsed interaction channels or cells that enter the interaction channels, (3) growth chambers with unoccupied regions, (4) abnormal cell growth that significantly alters the total number of cells in the growth chamber, or (5) cell growth near growth chambers may generate different diffusion rates. In all physically separated experiments (Experiments 1-6,10-13, **Table 1**), the connected chamber was excluded from the data set if a growth chamber was identified as an outlier based on these criteria (**Table 1**).

### Population-level fluorescence time-series analysis

Fluorescent time series measurements for each growth chamber in Experiments 5-6,10-13 (**Table 1**) were analyzed by bootstrapping, with p-values computed using bootstrap hypothesis testing. Using this method, the biological replicate curves for a given interaction channel length were randomly sampled 1000 times with replacement. In Experiment 1 (**Table 1**), background fluorescence was subtracted from the data by subtracting the minimum RFP fluorescence intensity across all growth chambers for model fitting (**Fig. 1c,d**).

In the forced oscillation experiments (Experiments 2-3, **Table 1**), a peak finding algorithm (Python) was applied to the time-series gene expression data at steady state with minimum inter-peak threshold of 21 min. The amplitude was computed by subtracting the minimum and maximum of each oscillation and dividing this value by two. To calculate the signal-to-noise ratio (SNR), a moving mean computed over 20 time points was subtracted from the data. The power spectra for each replicate was calculated using Welch’s method (Python) with a Hamming window applied across the length of the time-series. The power spectra were filtered to exclude frequencies lower than the signal bandwidth. The signal was defined as the total power of the signal bandwidth. The noise was computed as the total power of frequencies larger than the signal bandwidth. The power spectra for all the replicates for a given interaction channel length were randomly sampled with replacement 10,000 times. At each iteration, the signal-to-noise ratio was computed by dividing the signal by the noise. Bootstrap hypothesis testing was used to compute p-values.

For the dual-feedback oscillation experiment (Experiment 4, **Table 1**), the fluorescence intensity of each fluorescent reporter was normalized by subtracting the global minimum of the reporter across all replicates and dividing by the global maximum of fluorescence across all replicates. The mean-subtracted data was computed using a moving mean of 20 time points. A peak finding algorithm (Python) was applied to detect peaks with a minimum inter-peak distance of 70 min and a minimum peak height of 0.015 by analyzing the data after the media switch. The number of peaks detected, the amplitude of expression at each peak, and the distance between subsequent peaks were all computed for each replicate.

In the auxotroph experiments (Experiments 5, 6, 10-13, **Table 1**), the fluorescence background for each reporter was subtracted from the data and then the time-series was normalized by dividing by the maximum value. The change in fluorescence per unit time (ΔF Δt^−1^) was computed by determining the slope of a line fit to a 10 time point moving window and then multiplying by negative one. The global maximum of ΔF Δt^−1^ corresponded to the maximum growth rate. The doubling time was calculated using times the following equation:

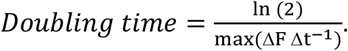

In Experiment 10 (**Table 1**), the ΔF Δt^−1^ curves displayed a biphasic trend characterized by a global and local maximum. To characterize the growth rate at each peak, the ΔF Δt^−1^ time-series was analyzed between the time point of the media switch and the time point corresponding to 25% of the maximum fluorescence. The local maxima within this time window were identified using the findpeaks algorithm (Python). The bootstrapped ΔF Δt^−1^ time-series were aligned by the first peak and the doubling times at the global maximum were calculated as described above. For the second growth phase, the doubling time was calculated at the maximum ΔF Δt^−1^ for the period of time between the global maximum and the time point corresponding to 25% of the maximum fluorescence.

### Single-cell image analysis

Single cell metrics were obtained with a custom machine learning approach implemented in Python with the Keras API running on top of TensorFlow^72^. We used two convolutional neural networks with U-Net architecture. First, we performed segmentation of individual cells in each image and then tracked each of the segmented cell instances over time. The segmentation network takes as an input the phase contrast images of cells grown in MISTiC and for each image and yields a binary mask segmenting the cells from the background. Training data was obtained from a separate experiment imaging fluorescently labeled *E. coli* at 60X magnification with phase contrast and fluorescence images collected every 10 minutes. We used the fluorescence images to generate binary segmentation masks of the cells, which then served as the ground truth for the phase contrast images used for network training. A total of 1066 images were curated this way. The network was trained for 100 epochs using a stochastic gradient descent (SGD) optimizer and a pixelwise weighted loss function to enforce the learning of narrow borders between adjacent cells. To minimize overfitting of the network to the training data, random affine transformations and elastic deformations were applied in real-time during the training process.

Cell tracking was performed with a separate U-Net similar to a method reported previously^73^. The input for this network is a set of consecutive binary segmentation masks. For each cell in the current segmentation, the network predicts the cell in the previous segmentation image from which the current cell was derived. This backwards tracking approach eliminates the need for the network to learn occurrences of cells leaving the chamber and reduces the number of classes to two (the tracked cell and the background). Using segmentations from the mixed auxotroph experiment, we curated 2656 sets of training images with a custom script in MATLAB. Training occurred for 200 epochs using an Adam optimizer and a class-weighted categorical cross-entropy loss function. Similarly, data augmentation was performed to reduce overfitting of the data.

Following segmentation and tracking, the raw output was processed with custom code in Python to reconstruct cell lineage and obtain single cell metrics. The instantaneous growth rate of each cell was computed from the cross-sectional area recorded during the 100-minute window (10 data points) immediately following that instant. Growth rate was computed by fitting a line to each 100 min window and then dividing the slope of the line by the average cell area during that time interval. For all analyses, a minimum tracking duration of 100 minutes was imposed to enforce consistent computation of growth rate. For all analyses involving growth rate, statistical outliers were identified using a modified z-score computed on the chamber averaged growth rates at each time point

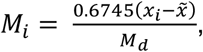

where 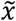 represents the median growth rate and *M*_*d*_ denotes the median absolute deviation^74^. Statistical outliers were detected using a threshold of *M*_*i*_ > 3.5. Growth chambers with more than one time point registering as an outlier were excluded from the analysis (**Table 1**). Experimental outliers occurred primarily due to segmentation and tracking errors caused by loss of focus or empty chambers at specific positions. Outliers were considered separately for each strain.

### Model fitting

Custom code (MATLAB) was used for computational modeling. An ordinary differential equation model was developed to study inter-strain communication via chemical signal diffusion (quorum-sensing). Detailed descriptions of the diffusion and gene expression models are in the Supplementary Information. The general mathematical form of the equations describing the concentration of AHL or fluorescein in each discretized spatial region is

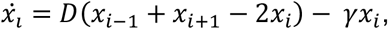

where *x*_*i*_ and *x*_*i+1*_ represent concentrations in adjacent regions of the device. The parameters *D* and *γ* denote the diffusion rate and degradation rate of the diffusible molecule, respectively. For the gene expression model, the general mathematical form for modeling transcription is

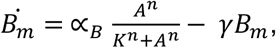

where *A* and *B*_*m*_ represent a transcription factor and its regulated transcript, respectively. The parameters α_*B*_, *n*, *K*, and *γ* denote the maximum transcription rate, Hill coefficient, half-maximum concentration or binding affinity and mRNA degradation rate, respectively. The general mathematical form for representing time delays due to sequential protein assembly, fluorescent protein maturation or media switching is

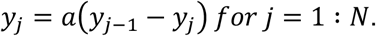

The species *y*_*N*_ represents the time-delayed species *y*_*1*_ and the delay time is computed by *N* · *a*^−1^.

The model was simulated using ode23s (MATLAB). A model with a variable number of delay equations was fit to the data using a genetic algorithm. The algorithm identified a best estimate for the parameter values and an optimal model structure by adjusting the number of delay equations to minimize the L^2^-norm between the model and the data. First, 100 parameter sets were randomly sampled using an upper and lower bound for each parameter. For each parameter set, the model was simulated and the L^2^-norm between the model and the data was computed. The parameters were ranked from lowest to highest L^2^-norm. The first parameter set (lowest L^2^-norm) was averaged with parameter sets 2-10, generating 9 new parameter sets. These parameter sets were combined with 81 randomly sampled parameter sets using an upper and lower bound for each parameter. This procedure was repeated until the L^2^-norm did not change significantly with additional iterations. The best estimates for the parameters are listed in **Table S3**.

The parameters of the amino-acid cross-feeding model were fit using a genetic algorithm. The genetic algorithm can be most efficient with high-order systems and many unknowns. One of the challenges with the genetic algorithm is there is no proof of convergence and the rate of convergence can be slow if the initial guesses on the parameters are far from the minimizing set and the bounds on the parameters are too broad. In order to overcome these challenges, careful consideration was taken into determining the lower and upper bounds on the parameters. Initially, bounds were determined based on biologically relevant and feasible values. Additionally, experimental observations where used to infer necessary relationships between parameters. The bounds on the parameters were adjusted accordingly. After this, the genetic algorithm was executed until the error became invariant for a sequence of 10 generations. Since the genetic algorithm is not optimal, it is possible to arrive at slightly different values if we were to run the genetic algorithm longer or reinitiate at new random initial conditions. However, the qualitative fits remain fairly close, as do the parameter values. Nevertheless, given experimental error, it is not in our benefit to achieve an optimal fit, since such a fit does not imply better prediction of quantitative values of parameters.

## ACKNOWLEDGEMENTS

We would like to thank John Crooks and Jeremy Schroeder for assistance with the laser pattern generator and Michael Chevalier, Ryan Clark, Bhuvana Krishnaswaymy, Freeman Lan, Austin McDaniel, Urbashi Mitra and Nimish Pujara and for helpful discussions. Research was sponsored by the Army Research Office and was accomplished under Grant Number W911NF-19-1-0269. S.G. was supported by NIH/NIGMS under award number T32 GM008293.

## AUTHOR CONTRIBUTIONS

O.S.V., P.A.R. and S.G. designed the research. S.G. and T.D.R. carried out experiments. S.G., M.G. and O.S.V. performed computational modeling. O.S.V., P.A.R., S.G. and T.D.R. discussed data analyses and S.G., O.S.V. and T.D.R. performed the analyses. O.S.V., S.G., P.A.R. and T.D.R. wrote the manuscript. J.L.G. and T.D.R. carried out the metabolite measurements. O.S.V. secured funding.

## DATA AVAILABILITY

The source data for Figures 1-5 is available as Source Data files.

## CODE AVAILABILITY

The code used for computational modeling and data analysis is available upon request.

