## Supplementary Information for "Investigating the dynamics of microbial consortia in spatially structured environments"

### Supplementary Information for
### Investigating the dynamics of microbial consortia in spatially structured environments

Sonali Gupta, Tyler D. Ross, Marcella M. Gomez, Job L. Grant, Philip A. Romero, Ophelia S. Venturelli

#### 1 Modeling diffusion through the interaction channel

We assumed that diffusion through the MISTiC interaction channels could be modeled as a one-dimensional process by discretizing space in the interaction channels into 1 micron regions. The chamber connected to the main channel with 10 $\mu$M fluorescein is the 'source' chamber, $x_1$, through which the dye enters the interaction channel and diffuses across at rate $D_1$. The concentration in the source chamber remains constant where

$$\dot{x_1} = 0. \tag{1}$$

We assumed that fluorescein degraded at a linear rate due to absorption by PDMS or photobleaching. The concentration of fluorescein in each region of the interaction channel is

$$\dot{x_l} = D_1(x_{l-1} + x_{l+1} - 2x_l) - \gamma_F x_l \text{ for } l = 2, 3, ..., L-1. \tag{2}$$

The 'sink' chamber, $x_L$ is on the opposite side of the interaction channel, through which fluorescein dilutes into the other main channel at a rate $D_2$. Fluorescein concentrations in the sink chamber are given by

$$\dot{x_L} = D_1(x_{L-1} - x_L) - D_2 x_L - \gamma_F x_L. \tag{3}$$

The model parameters (Table S1) were fit to the fluorescence data within the interaction channel using a nonlinear programming solver (fmincon, MATLAB). For $\gamma_F = 0$, the model exhibits linear chemical gradients that do not mirror the shape of the experimentally measured gradients (**Fig. S1**).

| **Parameter** | **Description** | **Value** |
|---|---|---|
| $D_1$ | Diffusion coefficient of fluorescein across the interaction channel | 2.0016 x $10^5$ time$^{-1}$ |
| $\gamma_F$ | Degradation rate of fluorescein | 7.4466 time$^{-1}$ |
| $F_c$ | Fluorescein concentration in source chamber | 0.9135 A.U. |
| $D_2$ | Diffusion coefficient of fluorescein into main channel | 1227.8 time$^{-1}$ |

Table S1: Estimated parameters for the fluorescein experiment.

### 2 Dynamic computational model of inter-strain communication across spatial distance

An ordinary differential equation model was developed to study quorum-sensing inter-strain communication across distance. Reflecting our diffusion model, we assumed that diffusion could be modeled as a one-dimensional process by discretizing space in the interaction channels into 1 micron regions. Dynamical delays were used to model sequential biochemical reactions, syringe activation, and transport of media from the inlets into the growth chambers. The model species are listed in Table S2.

| **Species** | **Description** |
|---|---|
| $w_i, i = 1, 2, ..., N_1$ | Arabinose concentration in different regions of the device |
| $GFP_m$ | GFP mRNA |
| $v_i, i = 1, ..., N_2$ | GFP protein intermediates |
| $v_{N_2}, GFP_p$ | Mature GFP protein |
| $LuxI_m$ | LuxI mRNA transcript |
| $u_i, i = 1, ..., N_3$ | LuxI protein intermediates |
| $u_{N_3}, LuxI_p$ | LuxI protein |
| $x_i, i = 1, 2, ..., L-1$ | AHL concentration in each region of the interaction channel |
| $x_L, AHL$ | AHL concentration within the receiver chamber |
| $LuxRAHL$ | Activated complex of $LuxR$ bound to $AHL$ |
| $RFP_m$ | RFP mRNA |
| $z_i, i = 1, ..., N_4$ | RFP protein intermediates |
| $z_{N_4}, RFP_p$ | RFP protein |

Table S2: Species descriptions for inter-strain communication model.

The following equations account for the time delay due to arabinose transport and diffusion into the device:

$$\dot{w_1} = a_1(ara - w_1), \tag{4}$$

$$\dot{w_i} = a_1(w_{i-1} - w_i) \text{ for } i = 2 : N_1 - 1, \tag{5}$$

$$\dot{w_{N_1}} = D_0(w_{N_1-1} - w_{N_1}). \tag{6}$$

The species $w_{N_1}$ represents the concentration of arabinose in the sender growth channel. The species $GFP_m$ is modeled as

$$G\dot{F}P_m = \alpha_{LuxI} \frac{w_{N_1}^{n_{LuxI}}}{K_{LuxI}^{n_{LuxI}} + w_{N_1}^{n_{LuxI}}} - \gamma_{GFP} GFP_m. \tag{7}$$

To account for time delays associated sequential assembly and maturation of GFP, the following delay equations were used:

$$\dot{v_1} = a_2(GFP_m - v_1), \tag{8}$$

$$\dot{v_j} = a_2(v_{j-1} - v_j) \text{ for } j = 2 : N_2. \tag{9}$$

The species $v_{N_2}$ represents concentration of mature GFP protein in the sender strain. The concentration of $LuxI_m$ was modeled as

$$Lu\dot{x}I_m = \alpha_{LuxI} \frac{w_{N_1}^{n_{LuxI}}}{K_{LuxI}^{n_{LuxI}} + w_{N_1}^{n_{LuxI}}} - \gamma_{LuxI} LuxI_m. \tag{10}$$

Similarly to GFP, we represent delays in translation of LuxI protein using the following equations:

$$\dot{u_1} = a_3(LuxI_m - u_1), \tag{11}$$

$$\dot{u_k} = a_3(u_{k-1} - u_k) \text{ for } k = 2 : N_3. \tag{12}$$

The species $u_{N_3}$ denotes the concentration of $LuxI_p$ in the sender strain. LuxI synthesizes AHL at a rate $p_{AHL}$, which diffuses into the interaction channel with a diffusion coefficient of $D_1$ and into the main channel with a diffusion coefficient of $D_2$. $D_2$ is significantly lower than $D_1$ since the diffusion rate through the cells is slower than the diffusion through the media. The equation representing AHL in the sender growth chamber is

$$\dot{x_1} = D_1(x_2 - x_1) - D_2 x_1 + p_{AHL} u_{N_3} - \gamma_{AHL} x_1. \tag{13}$$

The interaction chamber of length $L$ is discretized into approximately 1 $\mu m$ units, with $x_2$ and $x_{L-1}$ representing the region closest to the sender growth chamber or receiver growth chamber, respectively. The concentration of AHL in each region of the interaction channel is

$$\dot{x_l} = D_1(x_{l-1} + x_{l+1} - 2x_l) - \gamma_{AHL} x_l \text{ for } l = 2, 3, ..., L - 1. \tag{14}$$

The concentration of AHL in the receiver growth chamber is a balance between diffusion from the interaction channel and diffusion into the main channel and is modeled as

$$\dot{x_L} = D_1(x_{L-1} - x_L) - D_2 x_L - \gamma_{AHL} x_L. \tag{15}$$

In the receiver strain, we assume that the binding of AHL to LuxR is significantly faster than the timescales of transcription and translation. Therefore, based on the quasi-steady-state assumption, the concentration of activated LuxR is modeled as an algebraic equation

$$LuxRAHL = LuxR_{tot} \frac{x_L}{K_{LuxR} + x_L}. \tag{16}$$

The species $LuxRAHL$ activates the expression of $RFP_m$ using the following equation

$$R\dot{F}P_m = \alpha_{RFP} \frac{LuxRAHL^{n_{RFP}}}{K_{RFP}^{n_{RFP}} + LuxRAHL^{n_{RFP}}} - \gamma_{RFP} RFP_m. \tag{17}$$

We model time delays associated with translation and maturation of $RFP_p$ using the following equation

$$\dot{z_1} = a_4(RFP_m - z_1), \tag{18}$$

$$\dot{z_m} = a_4(z_{m-1} - z_m) \text{ for } m = 2 : N_4. \tag{19}$$

The species $z_{N_4}$ denotes the concentration of mature $RFP_p$ in the receiver strain.

### 3 Model of spatially separated amino-acid cross-feeding in microbial consortia

We constructed an ordinary differential equation model to study the effects of separation distance on amino acid cross-feeding dependent growth rates of strains in microbial communities. Similar to the quorum-sensing model, we assume that diffusion is modeled as a one-dimensional process by discretizing space in the interaction channels into one-micron regions. The diffusion coefficients of methionine (M) and phenylalanine (F) were set to the values of the inferred diffusion coefficients in the quorum-sensing model since these small molecules have similar molecular weights ($149 - 213$ $g\ mol^{-1}$).

| Parameter | Description | Value |
|---|---|---|
| $ara$ | Arabinose concentration | 0.0619 A.U. |
| $a_1$ | Time-varying rate coefficient | 1.3187 $\text{min}^{-1}$ |
| $N_1$ | Number of delay equations (arabinose) | 6 |
| $D_0$ | Diffusion coefficient into growth chamber | 0.057 $\text{min}^{-1}$ |
| $\alpha_{LuxI}$ | $LuxI_m$ maximum transcription rate | 264.53 A.U. $\text{min}^{-1}$ |
| $n_{LuxI}$ | LuxI Hill coefficient | 2.02 |
| $K_{LuxI}$ | $LuxI_m$ half-maximum concentration | 33.42 A.U. |
| $\gamma_{LuxI}$ | $LuxI_m$ degradation rate | 0.046 $\text{min}^{-1}$ |
| $N_3$ | Number of delay equations for LuxI translation & maturation | 6 |
| $\gamma_{GFP}$ | GFP mRNA degradation rate | 0.022 $\text{min}^{-1}$ |
| $a_2$ | Time-varying rate coefficient | 1.2690 $\text{min}^{-1}$ |
| $N_2$ | Number of delay equations for GFP translation and maturation | 15 |
| $a_3$ | Time-varying rate coefficient | 0.5607 $\text{min}^{-1}$ |
| $D_1$ | Diffusion coefficient for AHL diffusion into interaction channel | 7677.8 $\text{min}^{-1}$ |
| $D_2$ | Diffusion coefficient for AHL diffusion into main channels | 251.0 $\text{min}^{-1}$ |
| $p_{AHL}$ | AHL production rate | 564.5 $\text{min}^{-1}$ |
| $\gamma_{AHL}$ | AHL degradation rate | 0.086 $\text{min}^{-1}$ |
| $LuxR_{tot}$ | Total concentration of LuxR protein | 9.5 A.U. |
| $K_{LuxR}$ | Half-maximum concentration for $LuxRAHL$ | 54.2 A.U. |
| $\alpha_{RFP}$ | Maximum transcription rate for $RFP_m$ | 184 A.U. $\text{min}^{-1}$ |
| $n_{RFP}$ | Hill coefficient for RFP transcription | 1.04 |
| $K_{RFP}$ | Half-maximum concentration for $RFP_m$ | 101.3 A.U. |
| $\gamma_{RFP}$ | $RFP_m$ degradation rate | 0.037 $\text{min}^{-1}$ |
| $a_4$ | Time-varying rate coefficient | 0.9195 $\text{min}^{-1}$ |
| $N_4$ | Number of delay equations for RFP translation/maturation | 4 |

Table S3: Parameter values for quorum-sensing model. Time delays can be computed as the ratio of the number of delay equations to the time-varying rate coefficient. This yields 6.0 min for arabinose transport from the inlets into the sender growth chambers, 11.8 min for GFP translation and maturation and 4.4 min for RFP translation and maturation.

| Species | Description |
|---|---|
| $[M]_{\Delta metA}$ | Intracellular [M] in $\Delta metA$ |
| $[F]_{\Delta pheA}$ | Intracellular [F] in $\Delta pheA$ |
| $x_{1,\ldots,L}, [M]_{ext}$ | Extracellular [M] in different regions of the interaction channel or growth chambers |
| $y_{1,\ldots,L}, [F]_{ext}$ | Extracellular [F] in different regions of the interaction channel or growth chambers |

Table S4: Species descriptions for auxotroph model.

#### 3.1 Intracellular transport of amino acids from the extracellular environment

We compute the intracellular concentrations of M and F based on the extracellular concentration using a previous method (Perez AM et al. *Molecular Systems Biology* 2017). The intracellular concentration of M in the $\Delta metA$ strain is given by

$$[M]_{\Delta metA} = U_M \frac{[M]_{ext}}{\kappa_M + [M]_{ext}} + [M]_{ext}, \tag{20}$$

where

$$U_M = \frac{V_{max,M} P_{tot}}{D_c} \quad \text{and} \quad \kappa_M = \frac{V_{max,M}}{k_M}. \tag{21}$$

Here, $[M]_{ext}$ is the concentration of extracellular methionine, $D_c$ is the diffusion coefficient of amino acid across the outer membrane, $V_{max,M}$ is the uptake rate of M, $P_{tot}$ is the concentration of membrane transporters, and $k_M$ is the binding affinity of M to the transporter. Similarly, the concentration of F in the $\Delta pheA$ strain is

$$[F]_{\Delta pheA} = U_F \frac{[F]_{ext}}{\kappa_F + [F]_{ext}} + [F]_{ext}, \tag{22}$$

where

$$U_F = \frac{V_{max,F} P_{tot}}{D_c} \quad \text{and} \quad \kappa_F = \frac{V_{max,F}}{k_F}. \tag{23}$$

Here, $[F]_{ext}$ denotes the extracellular F concentration, $V_{max,F}$ is the uptake rate of M, and $k_F$ is the binding affinity of F to its transporter. We assume that $D_c$ and $P_{tot}$ are equivalent for both strains. The kinetic parameters $V_{max,M}$, $V_{max,F}$, $\kappa_M$, and $\kappa_F$ were derived from Piperno JR and Oxender DL *Journal of Biological Chemistry* 1968 (**Table S5**).

#### 3.2 Amino acid limited growth rates of auxotroph strains

We assume that growth of each strain is limited by the concentration of the amino acid that this strain is deficient to produce. Therefore, the instantaneous normalized growth rate of each strain is a function of the intracellular concentration of the amino acid. We assume a different non-zero basal growth rate for $\Delta metA$ and $\Delta pheA$ in the absence of the rescuing amino acid denoted by $r_M$ and $r_F$, respectively. The normalized growth rates for $\Delta metA$ and $\Delta pheA$ are given by the equations

$$GR_{\Delta metA} = (1 - r_M) \frac{[M]^{b_M}_{\Delta metA}}{M^{b_M}_{EC50} + [M]^{b_M}_{\Delta metA}} + r_M, \tag{24}$$

$$GR_{\Delta pheA} = (1 - r_F) \frac{[F]^{b_F}_{\Delta metA}}{F^{b_F}_{EC50} + [F]^{b_F}_{\Delta pheA}} + r_F. \tag{25}$$

The parameters $M_{EC50}$ and $F_{EC50}$ represent the concentrations of M and F required by $\Delta metA$ and $\Delta pheA$, respectively, to achieve a half-maximal growth rate. The parameters $b_M$ and $b_F$ are Hill coefficients that map the concentration of the amino acid to the growth rate of each strain.

#### 3.3 Production and diffusion of amino acids

We model the production and diffusion of the amino acids M and F between growth chambers. The $\Delta pheA$ and $\Delta metA$ strains are located in the left and right growth chambers and produce M and F that diffuse through the interaction channel to the opposite end of the channel. The strains $\Delta pheA$ or $\Delta metA$ release and consume M or F. We assume that the amino acid consumption rate cannot be larger than its production rate. The maximum net production of M and F is represented by $\alpha_{net,M}$ and $\alpha_{net,F}$ and is defined as the maximum difference between the production rate minus the consumption rate of each amino acid. We model the net production, degradation, and diffusion of M and F. Let $x_i = [M]$ for $i = 1, ..., L$ denote the concentration of M in the $i^{th}$ unit of the channel.

$$\dot{x}_1 = D_1(x_2 - x_1) - D_2 x_1 - \gamma_M x_1 + \alpha_M \frac{(GR_{\Delta pheA}/U_F)^{c_M}}{(GR_{\Delta pheA}/U_F)^{c_M} + 1} \tag{26}$$

$$\dot{x}_i = D_1(x_{i-1} + x_{i+1} - 2x_i) - \gamma_M x_i \qquad \text{for } i = 2, 3, \ldots, L-1 \tag{27}$$

$$\dot{x}_L = D_1(x_{L-1} - x_L) - D_2 x_L - \gamma_M x_L \tag{28}$$

$$\tag{29}$$

Similarly, let $y_i = [F]$ for $i = 1, ..., L$ denote the concentration of F in the $i^{th}$ unit of the channel.

$$\dot{y}_1 = D(y_2 - y_1) - D_2 y_1 - \gamma_F y_1 \tag{30}$$

$$\dot{y}_k = D(y_{i-1} + y_{i+1} - 2y_i) - \gamma_F y_i \qquad \text{for } i = 2, 3, \ldots, L-1 \tag{31}$$

$$\dot{y}_L = D(y_{L-1} - y_L) - D_2 y_L - \gamma_F y_L + \alpha_F \frac{1}{((GR_{\Delta metA}/U_M)^{c_F} + 1)} \tag{32}$$

$$\tag{33}$$

Note that $[M]_{ext} = x_L$ and $[F]_{ext} = y_1$. The parameters $\gamma_M$ and $\gamma_F$ denote degradation rates of M and F, $\alpha_M$ and $\alpha_F$ are maximum production rates of the amino acids, and $c_M$ and $c_F$ are Hill coefficients for amino acid production. Consistent with our metabolite measurements, the net production of M is proportional to the growth rate of $\Delta pheA$, whereas the net production of F is inversely proportional to the growth rate of $\Delta metA$.

| Parameter | Description | Value |
|---|---|---|
| $\alpha_M$ | Maximum production of M by $\Delta pheA$ | 0.045 mM $\text{min}^{-1}$ |
| $\alpha_F$ | Maximum production of F by $\Delta metA$ | 0.55 mM $\text{min}^{-1}$ |
| $c_M$ | Hill coefficient of M production | 1.35 |
| $c_F$ | Hill coefficient of F production | 1.63 |
| $U_M$ | Maximum uptake rate of M by $\Delta metA$ | 0.022 $\text{min}^{-1}$ |
| $U_F$ | Maximum uptake rate of F by $\Delta pheA$ | 4.79 $\text{min}^{-1}$ |
| $M_{EC50}$ | [M] that enables half-maximum growth of $\Delta metA$ | 0.35 mM |
| $F_{EC50}$ | [F] that enables half-maximum growth of $\Delta pheA$ | 0.12 mM |
| $r_M$ | Basal growth rate of $\Delta metA$ in absence of M | 0.073 $\text{min}^{-1}$ |
| $r_F$ | Basal growth rate of $\Delta pheA$ in absence of F | 0.21 $\text{min}^{-1}$ |
| $\gamma_M$ | Degradation rate of methionine | 0.073 $\text{min}^{-1}$ |
| $\gamma_F$ | Degradation rate of phenylalanine | 0.044 $\text{min}^{-1}$ |
| $b_M$ | Hill coefficient of $\Delta metA$ growth | 1.81 |
| $b_F$ | Hill coefficient of $\Delta pheA$ growth | 4.06 |
| $D_1^*$ | Diffusion coefficient through interaction channel | 7677.8 $\mu$m $\text{min}^{-1}$ |
| $D_2^*$ | Diffusion coefficient from growth chambers to main channels | 251.0 $\mu$m $\text{min}^{-1}$ |
| $V^+_{max,M}$ | Maximum uptake rate of M | 0.39 |
| $V^+_{max,F}$ | Maximum uptake rate of F | 0.75 |
| $\kappa^+_M$ | Ratio of uptake rate to transporter binding affinity of M | 0.0023 mM |
| $\kappa^+_F$ | Ratio of uptake rate to transporter binding affinity of F | 0.00072 mM |
| $P_{tot}/D_c$ | Diffusion coefficient of amino acid across cell membranes | 1 |

Table S5: Parameter values for auxotroph model. The symbols * and + indicates parameters from the quorum-sensing model or previous literature (Piperno JR and Oxender DL *Journal of Biological Chemistry* 1968), respectively.

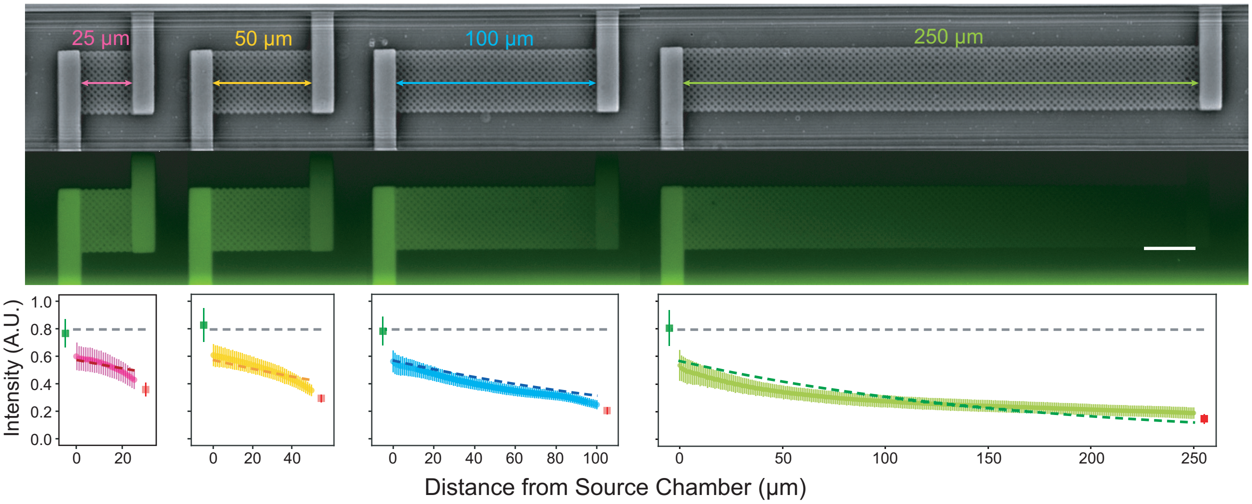


Figure S1: **Fluorescein dye demonstrates chemical gradients established within MISTiC. Top:** Phase contrast and fluorescence microscopy images of the microfluidic device. Fluorescein and water were continuously administered through separate main channels. The scale bar represents 25 $\mu m$. **Bottom:** fluorescence intensity in the growth chambers and interaction channels as a function of distance from the source chamber. Green and red squares represent the average intensity within source and sink chambers, respectively. The difference in the average fluorescence intensity between the 25 $\mu m$ and 250 $\mu m$ source channel was not statistically significant (P = 0.7539) based on an unpaired t-test. The difference in the average fluorescence intensity between the 25 $\mu m$ and 250 $\mu m$ sink channels was statistically significant (P = 0.0083) based on an unpaired t-test. The grey dashed line represents the mean fluorescence intensity in the source chambers across the different interaction channel lengths. Dashed lines represent the model fit to the data. Data points represent average fluorescein concentration in 1 $\mu m$ regions of the interaction channels. Error bars represent one standard deviation from the mean of three interaction channels in different regions of the device.

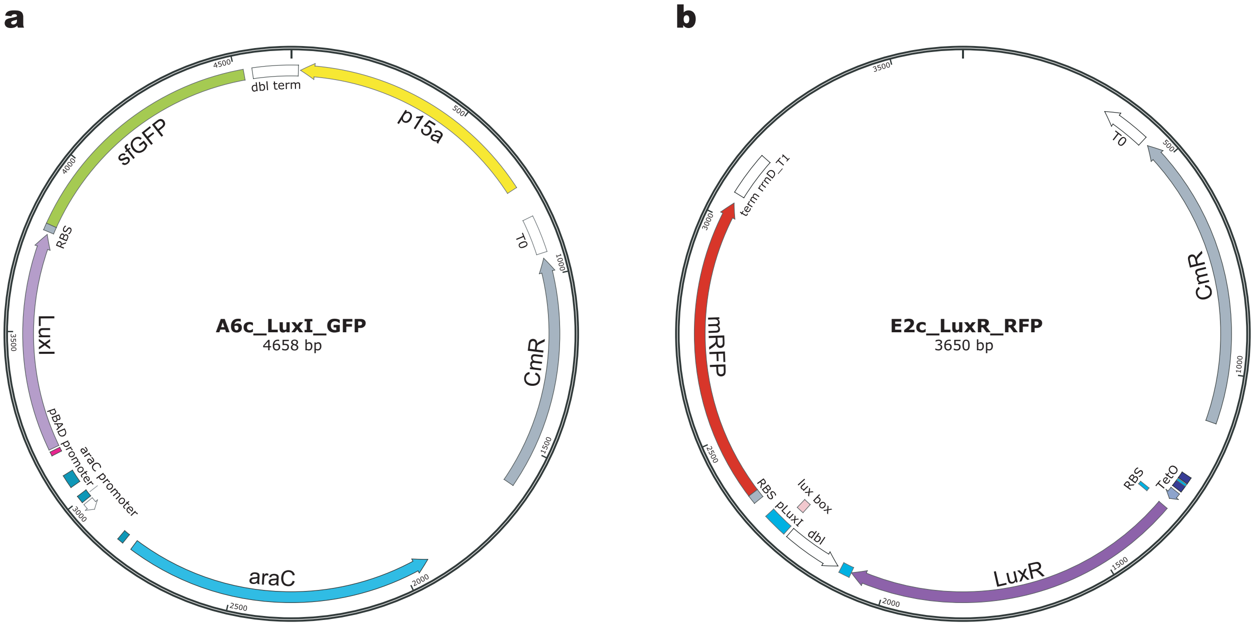


Figure S2: **Plasmid maps for sender-receiver quorum sensing experiments.** **(a)** Plasmid map for A6c_LuxI_GFP, which was transformed into the sender *E. coli* strain BW27783 (Materials and Methods). The arabinose-inducible $P_{BAD}$ promoter activates transcription of the operon containing the synthetase LuxI and fluorescent reporter GFP. **(b)** Plasmid map for E2c_LuxR_RFP that was transformed into the receiver *E. coli* strain MG1655z1 (Materials and Methods). The key features include the receptor LuxR regulated by the aTc-inducible $P_{TET}$ promoter and the fluorescent reporter RFP driven by a $P_{LuxI}$ promoter.

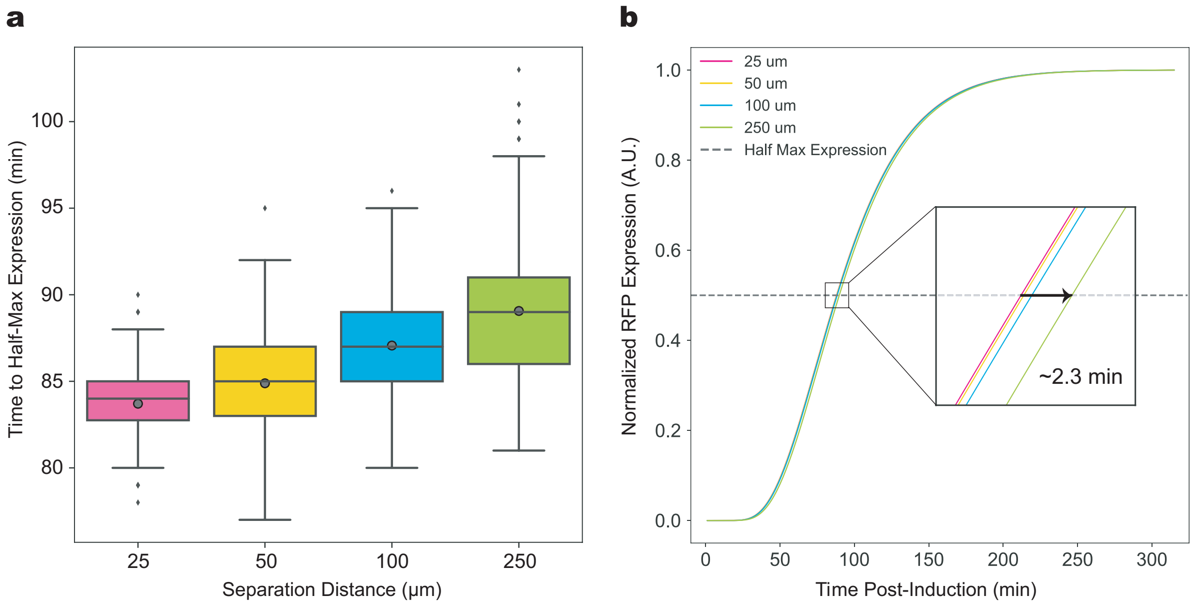


Figure S3: **Characterization of time delays in quorum-sensing chemical signal communication. (a)** Experimentally measured time delays for the quorum-sensing synthetic community as a function of the distance separating the sender and receiver strains. The distributions were computed by bootstrapping the RFP fluorescence as a function of time (1000 resampled curves) (Materials and Methods). For each resampled curve, the time delay was calculated by normalizing the curve between 0 and 1 and determining the time to reach 0.5. Lines and circles within the boxes represent the median and mean of the distributions, respectively. Upper and lower box edges represent upper and lower quartiles, respectively. Upper and lower whiskers represent the 95th and 5th confidence intervals, respectively. **(b)** Normalized model simulations of $RFP_p$ over time for different distances from the sender strain. Increasing the distance from 25 $\mu m$ to 250 $\mu m$ resulted in a 2.3 min time delay computed as the time to reach the half-maximum $RFP_p$ concentration (dashed line).

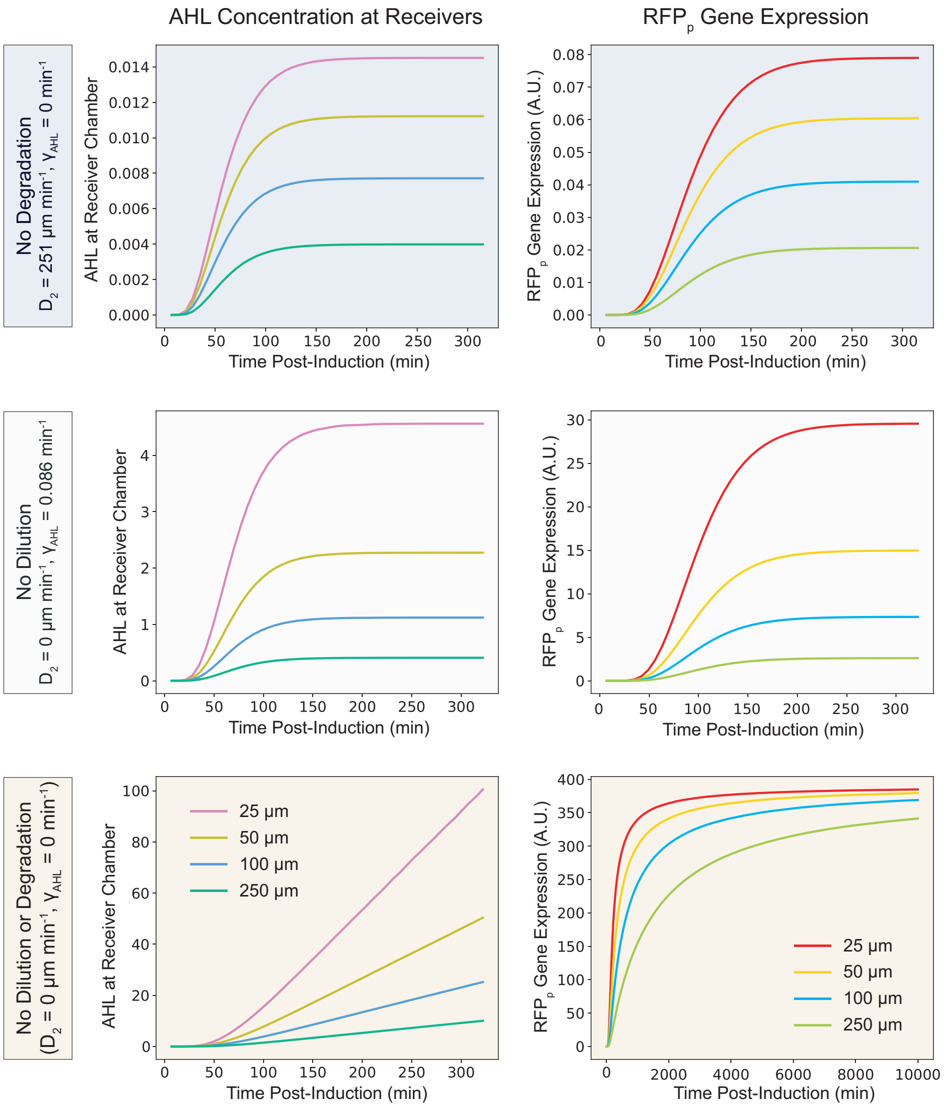


Figure S4: **Parameter dependence of simulated** $AHL$ **and** $RFP_p$**. Top row:** model time-responses of $AHL$ (left column) and $RFP_p$ (right column) where the degradation rate of AHL in the interaction channel is set to zero ($\gamma_{AHL} = 0$). **Middle row:** model time-responses where the diffusion constant of $AHL$ away from the growth chambers into the main channel is set to zero ($D_2 = 0$). **Bottom row:** model time-responses of $AHL$ and $RFP_p$ where $D_2 = 0$ and $\gamma_{AHL} = 0$.

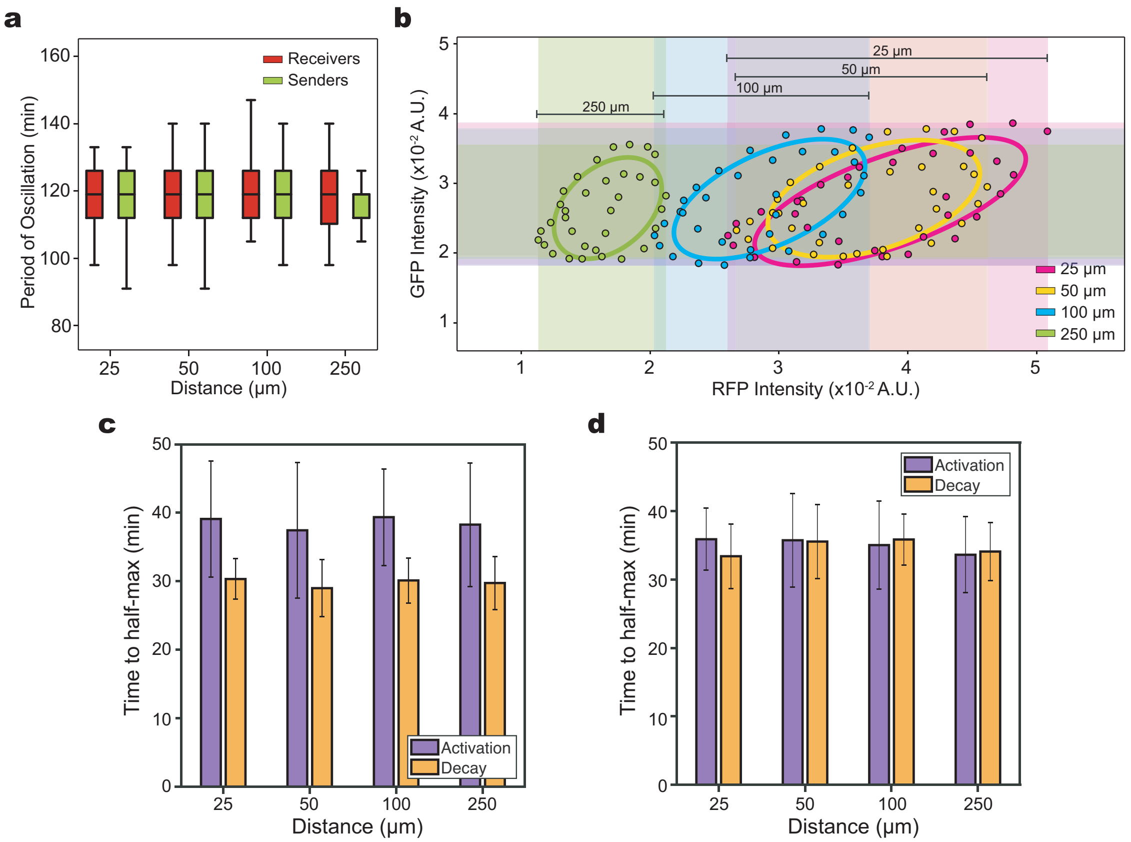


Figure S5: **Effects of spatial separation on the response of inter-strain communication to a periodic input.** **(a)** Box plots of the experimentally measured distribution of oscillatory periods, determined by applying a peak finding algorithm to time-series RFP (receivers) or GFP (senders) data for all cell growth chambers (Experiment 2, **Table 1**). The period was measured as a two hour oscillation (defined as 1 hr exposed to arabinose and 1 hr in the absence of arabinose) between 304 to 1004 min. Horizontal lines within boxes indicate the median and upper and lower box edges represent the upper and lower quartiles, respectively. Upper and lower whiskers represent the 95th and 5th confidence intervals, respectively. **(b)** Phase plot of RFP vs. GFP for two oscillatory periods from 580 to 700 min for a forced oscillation experiment with an arabinose period of 2 hr (Experiment 2, **Table 1**). **(c)** Activation and decay response times for GFP (sender strain). Activation response time is defined as the time to reach half-maximum from the minimum to the maximum of each period. The decay response time is defined as the time to reach half-minimum from the maximum to the minimum of each period. Error bars represent one standard deviation from the mean. **(d)** Activation and decay response times for RFP (receiver strain). Error bars represent one standard deviation from the mean.

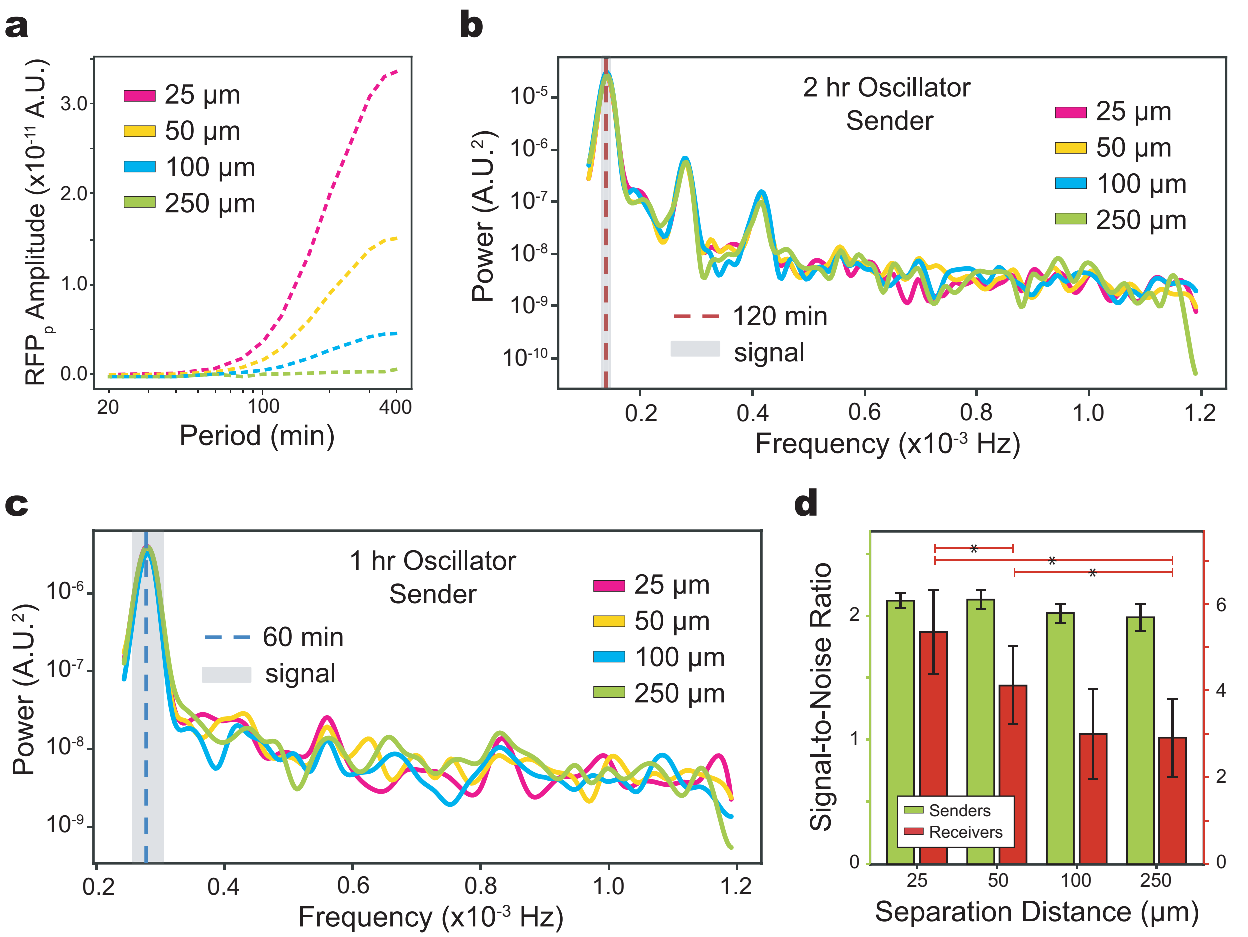


Figure S6: **Amplitude variation, power spectra, and signal-to-noise ratios for the sender-receiver periodic input experiments.** **(a)** Relationship between the period of an oscillatory arabinose ($ara$) input and the $RFP_p$ steady-state amplitude in the model for $n_{RFP} = 3$ for different distances from the sender strain. **(b)** Power spectrum (PS) of GFP (sender strain) in response to a periodic input with a period of 2 hr (Experiment 2, **Table 1**). PS was calculated using the Welch method with a Hamming windowing function at steady-state after 395 min. The arabinose frequency is indicated with a dashed red line. The signal bandwidth, denoted by a gray shaded region, is defined as the frequency range corresponding to 115-125 min. **(c)** PS for GFP (sender strain) for the forced oscillation experiment input period of 1 hr (Experiment 3, **Table 1**). PS was calculated using the Welch method with a Hamming windowing function at steady-state after 500 min. The arabinose frequency is represented by a dashed red line. The signal bandwidth is represented by a gray shaded region and is defined as the frequency range corresponding to 55-65 min. **(d)** Signal-to-noise (SNR) ratios based on the PS of GFP or RFP (receiver strain) across distance for the forced oscillation experiment with an input period of 2 hr (Experiment 2, **Table 1**). Left and right axes denote SNR for GFP and RFP, respectively. Error bars represent one standard deviation from the mean. Horizontal lines with stars (colored green for senders and red for receivers) denote a statistically significant difference ($P < 0.05$) based on bootstrapped hypothesis testing.

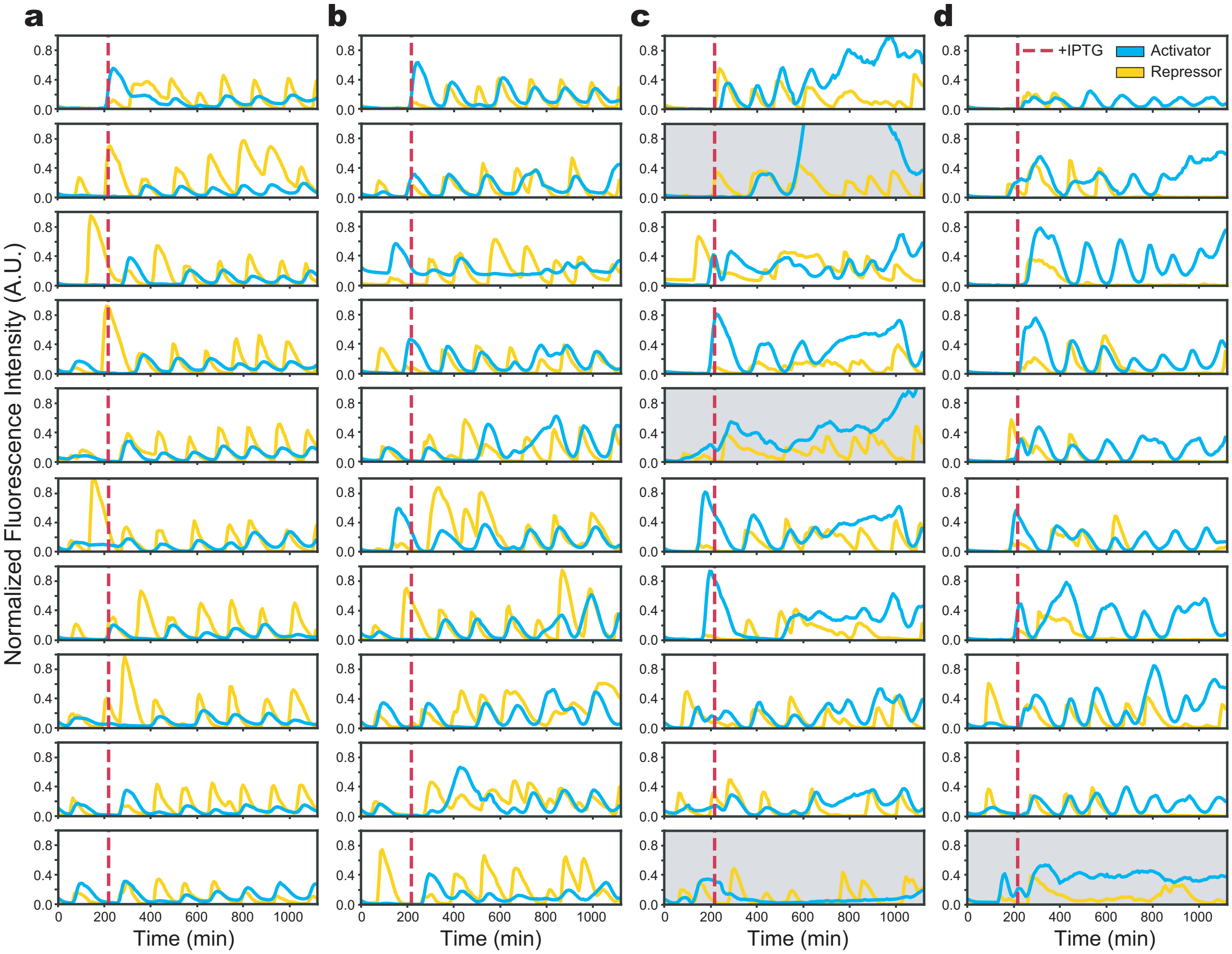


Figure S7: **Fluorescence intensity over time of dual-feedback oscillators in the presence of 1 mM IPTG** (Experiment 4, **Table 1**). **(a)** Normalized CFP (activator) and YFP (repressor) fluorescence intensity as a function of time for a separation distance of 25 $\mu m$. The red dashed line indicates the time when IPTG was introduced. For each fluorescence reporter, the fluorescence intensity was normalized between the global minimum and maximum excluding the outliers (gray shaded boxes). **(b)** Normalized CFP (activator) and YFP (repressor) fluorescence intensity as a function of time for a separation distance of 50 $\mu m$. **(c)** Normalized CFP (activator) and YFP (repressor) fluorescence intensity as a function of time for a separation distance of 100 $\mu m$. **(d)** Normalized CFP (activator) and YFP (repressor) fluorescence intensity as a function of time for a separation distance of 250 $\mu m$.

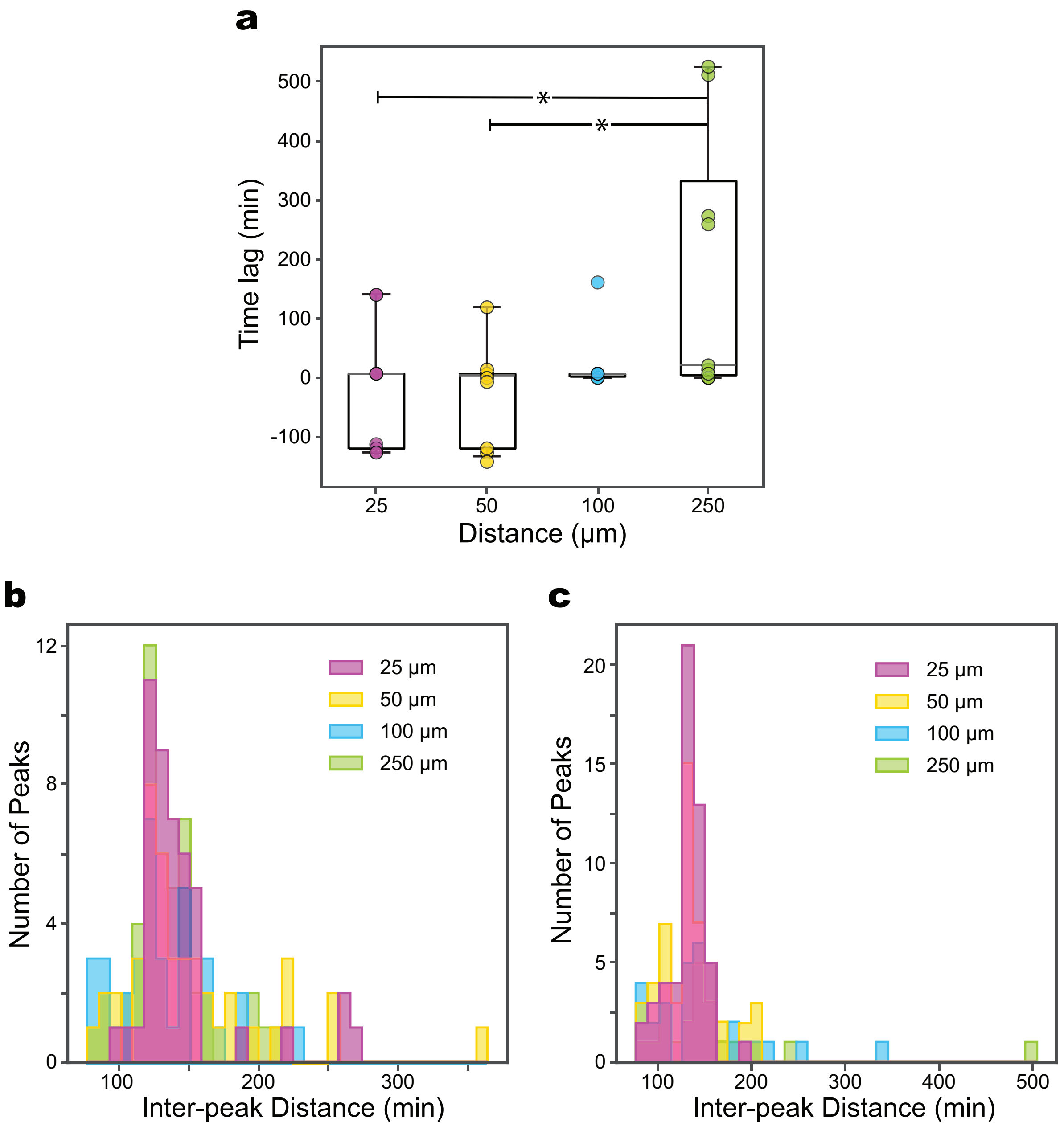


Figure S8: **Inter-peak distances and time lags of maximum cross-correlation for the dual-feedback oscillator consortium** (Experiment 4, **Table 1**). **(a)** Box plot denoting time lags of maximum cross-correlation between CFP (activator) and YFP (repressor) for paired interaction channels. Horizontal lines within boxes denote the median and upper and lower edges represent the upper and lower quartiles, respectively. Upper and lower whiskers represent the 95th and 5th confidence intervals, respectively. The horizontal bar with a star (*) indicates a statistically significant difference ($P < 0.05$) based on an unpaired t-test (**Fig. S14**). **(b)** Histogram of inter-peak distances for CFP (activator). A peak finding algorithm (Python) was used to determine the peaks in the normalized CFP fluorescence intensity as a function of time. CFP was normalized by subtracting a moving mean. The inter-peak distance was computed difference in the times corresponding to each peak for all replicates of a given interaction channel length. **(c)** Histogram of inter-peak distances for YFP (repressor).

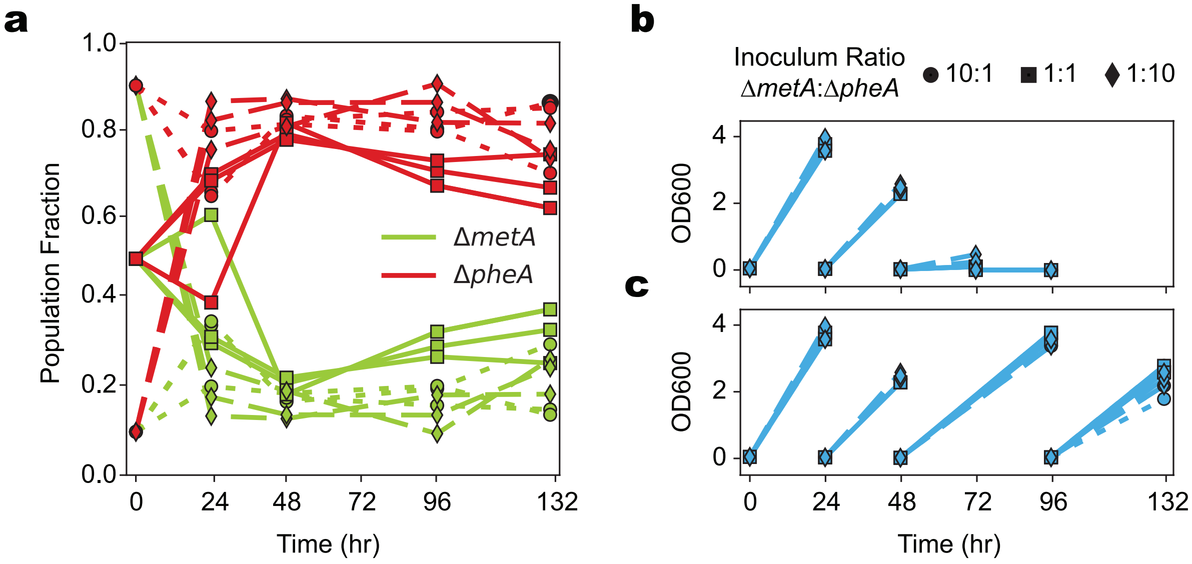


Figure S9: **Population dynamics of the Δ*metA*-Δ*pheA* consortium in batch culture**. **(a)** Fraction of each strain as a function of time. Marker and line styles correspond to different initial ratios of Δ*metA* to Δ*pheA*. Circles and dotted lines represent 10:1, squares and solid lines represent 1:1, and diamonds and dashed lines represent 1:10. **(b)** OD600 measurements as a function of time for a passaging period maintained at 24 hours. **(c)** OD600 measurements as a function of time for a second condition where the incubation period after the second passage was extended to 48 hr. The number of cells counted for each condition ranged between 250-3960.

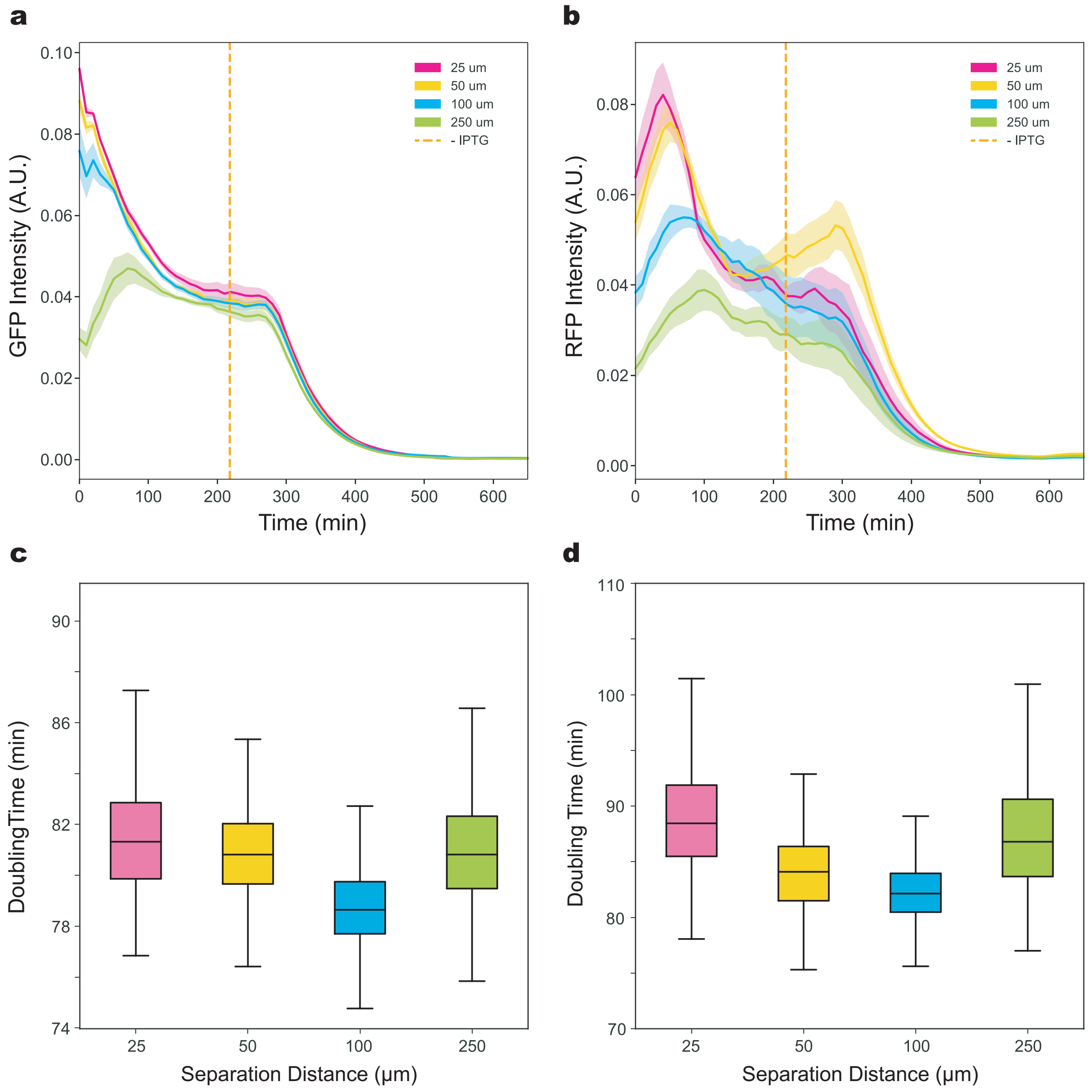


Figure S10: **Fluorescence as a function of time and doubling time distributions for the +M +F condition** (Experiment 5, **Table 1**). **(a)** GFP fluorescence in the $\Delta metA$ strain as a function of time in media supplemented with all amino acids. The dashed line indicates the time of the media switch from media containing IPTG to media lacking IPTG. Solid lines and shaded regions represent the mean and one standard deviation from the mean, respectively. **(b)** RFP fluorescence in the $\Delta pheA$ strain over time in media supplemented with all amino acids. The dashed line represents the media switch for removal of IPTG. **(c)** Box plot of cell doubling times for $\Delta metA$ at each interaction channel length. Horizontal lines within boxes depict the median and upper and lower edges represent the upper and lower quartiles, respectively. Upper and lower whiskers represent the 95th and 5th confidence intervals, respectively. **(d)** Box plot of the $\Delta pheA$ doubling times at each interaction channel length.

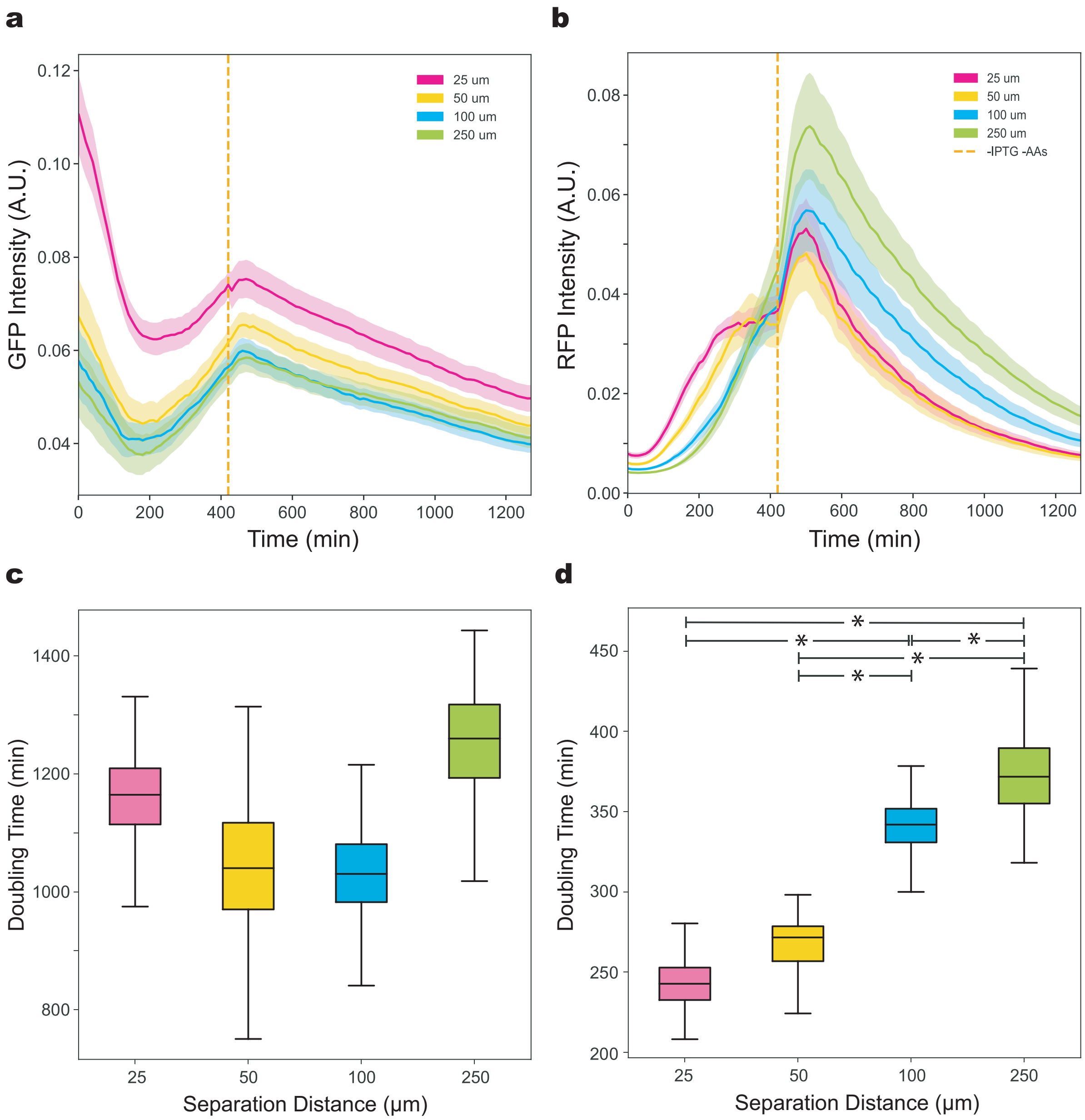


Figure S11: **Fluorescence as a function of time and doubling time distributions for the -M-F condition** (Experiment 6, **Table 1**). **(a)** GFP fluorescence over time in the $\Delta metA$ strain at different distances from the partner strain ($\Delta pheA$). The dashed line represents the time when the media was switched from pre-culture to test conditions. Solid lines and shaded regions represent the mean and one standard deviation from the mean, respectively. **(b)** RFP fluorescence over time in the $\Delta pheA$ strain at different distances from the partner strain ($\Delta metA$). **(c)** Box plot of the cell doubling times for the $\Delta metA$ strain. Horizontal lines within boxes denote the median and upper and lower edges represent the upper and lower quartiles, respectively. Upper and lower whiskers represent the 95th and 5th confidence intervals, respectively. **(d)** Box plot of the doubling times for $\Delta pheA$ strain. The horizontal bar with stars (*) indicates a statistically significant difference ($P < 0.05$) based on an unpaired t-test (**Fig. S15**).

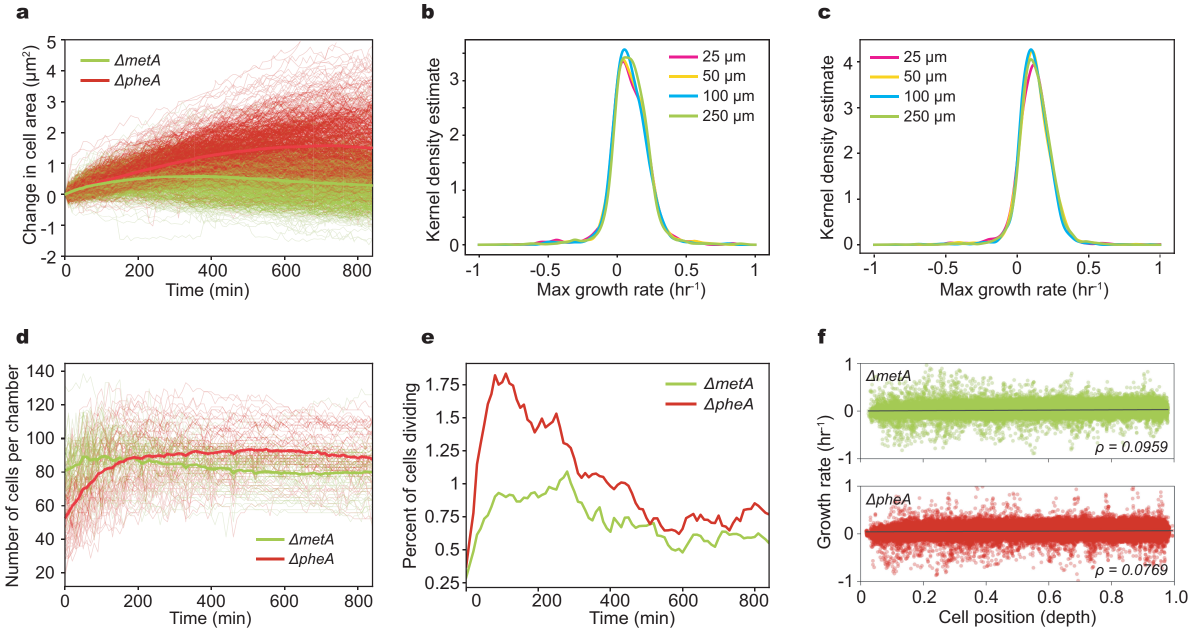


Figure S12: **Quantification of single-cell growth in the mixed auxotroph condition** (Experiment 7, **Table 1**). **(a)** Change in cell area as a function of time for $\Delta metA$ (n = 937) and $\Delta pheA$ (n = 662). The change in cell area is defined as the cumulative change in cross-sectional area of each cell from its initial value at the time of the media switch. Thin and bold lines represent individual cells and the strain averages, respectively. **(b)** Kernel density estimate distribution of $\Delta metA$ single cell growth rates over 14 hr post media switch in the mixed auxotroph experiment for each interaction channel length. **(c)** Kernel density estimate distribution of $\Delta pheA$ single cell growth rates over 14 hr post media switch in the mixed auxotroph experiment for each interaction channel length. **(d)** Total number of $\Delta metA$ and $\Delta pheA$ cells in each growth chamber as a function of time. Bold lines represent the average number of cells for each strain across all growth chambers. **(e)** Percent of $\Delta metA$ or $\Delta pheA$ cells dividing as a function of time. The fraction of dividing cells was computed using a window spanning 60 min starting at each time point, which was then divided by the average number of $\Delta metA$ or $\Delta pheA$ cells in the window. **(f)** Scatter plot of the correlation between the growth rate and depth position of individual $\Delta metA$ (n=165408) and $\Delta pheA$ (n=193970) cells in a mixed community (Experiment 7, **Table 1**). Positions zero and one represent the outermost and deepest regions of the growth chamber, respectively, relative to the main channels. Solid bold lines show simple linear regression between maximum growth rate and chamber position. The Pearson correlation coefficient is 0.0959 ($P < 0.05$) for $\Delta metA$ and 0.0769 ($P < 0.05$) for $\Delta pheA$.

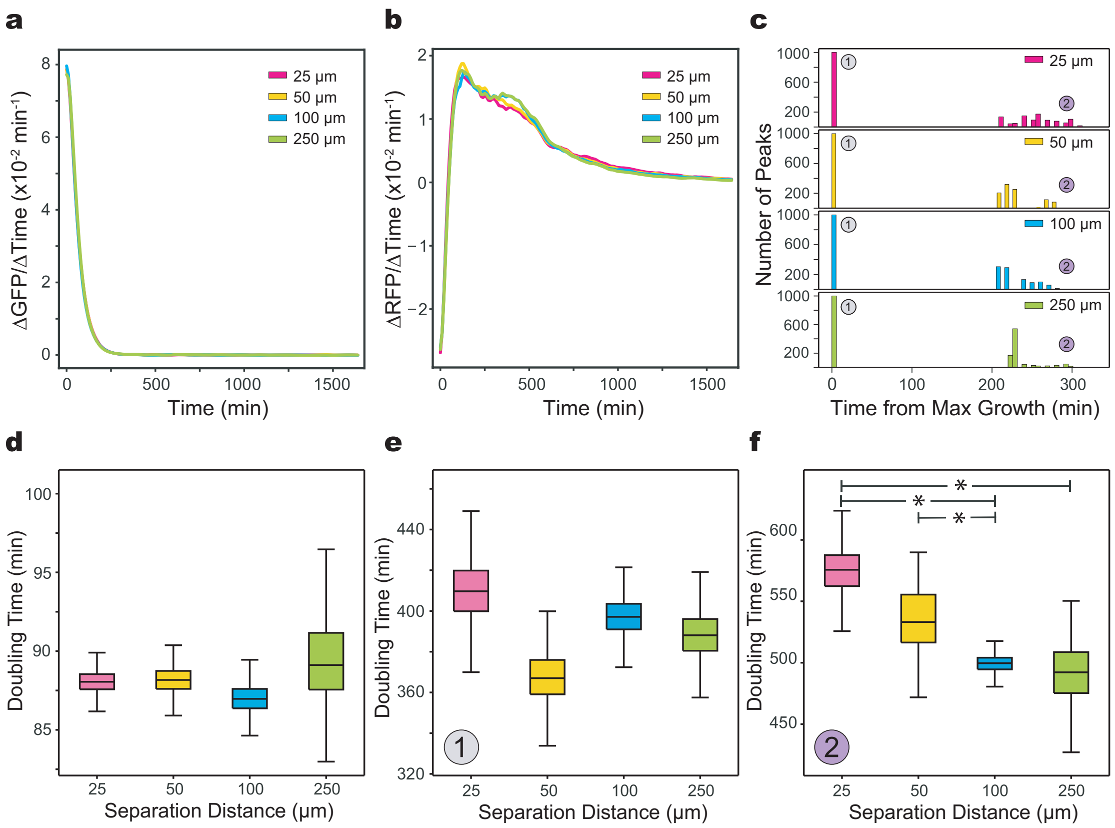


Figure S13: **Characterization of population-level growth rates of amino acid auxotrophs in the +M-F condition** (Experiment 8, **Table 1**). **(a)** Rate of change in GFP fluorescence of Δ*metA* as a function of time. Solid lines and shaded regions represent the mean and one standard deviation from the mean, respectively. **(b)** Rate of change of RFP fluorescence of Δ*pheA* over time. **(c)** Histogram of the number of peaks identified in the rate of change of RFP fluorescence over time for the 1000 bootstrapped curves using a peak finding algorithm (Materials and Methods). The distributions were normalized to the time of maximum growth rate. **(d)** Box plot of cell doubling times for the Δ*metA* strain at different distances from the Δ*pheA* strain. Horizontal lines within boxes depict the median and upper and lower edges represent upper and lower quartiles, respectively. Upper and lower whiskers represent the 95th and 5th confidence intervals, respectively. **(e)** Box plot of cell doubling times for first growth phase corresponding to the maximum growth rate of the Δ*pheA* strain. **(f)** Box plot of the cell doubling times for second growth phase of the Δ*pheA* strain. The horizontal bars denote a statistical significant difference ($P < 0.05$) based on an unpaired t-text (**Fig. S15**).

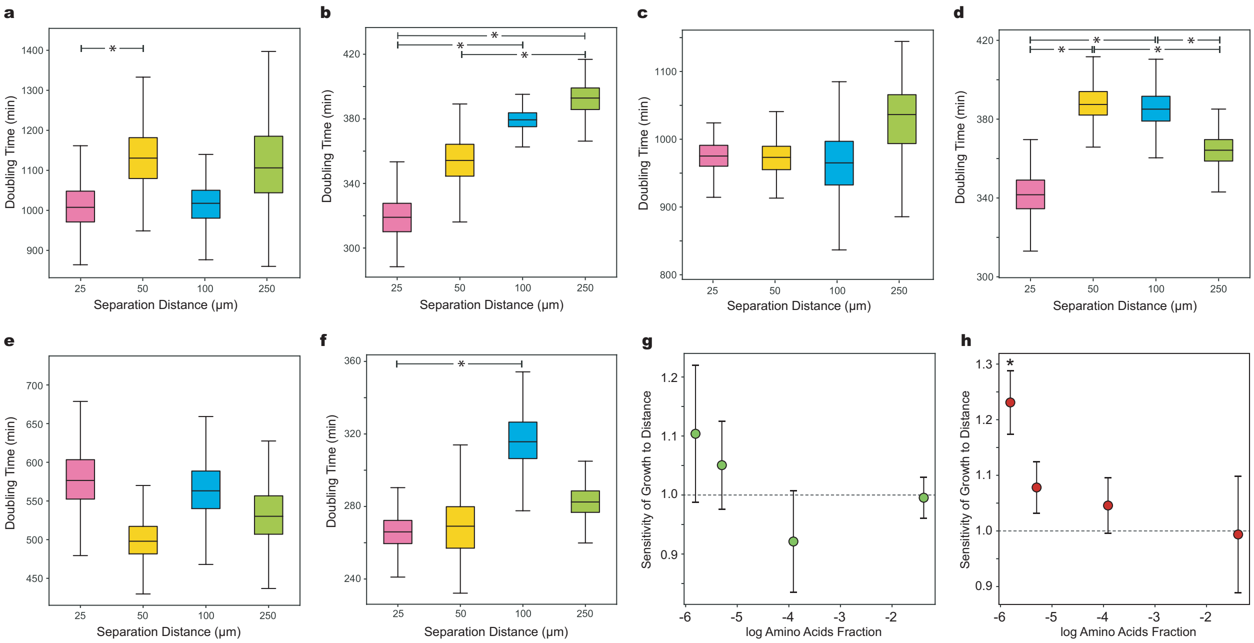


Figure S14: **Population-level growth rates of amino-acid auxotrophs across different concentrations of supplemented amino acids.** **(a)** Box plot of the doubling times for the $\Delta metA$ strain in Experiment 9 (**Table 1**). For each box, horizontal lines depict the median and upper and lower box edges represent the upper and lower quartiles, respectively. Upper and lower whiskers represent the 95th and 5th confidence intervals, respectively. The horizontal bars denote a statistically significance difference (P < 0.05) based on an unpaired t-test (**Fig. S15**). **(b)** Box plot of the cell doubling times for the $\Delta pheA$ strain for Experiment 9. **(c)** Box plot of the cell doubling times for the $\Delta metA$ strain in Experiment 10. **(d)** Box plot of the cell doubling times for the $\Delta pheA$ strain in Experiment 10. **(e)** Box plot of the cell doubling times for the $\Delta metA$ strain in Experiment 11. **(f)** Box plot of the cell doubling times for the $\Delta pheA$ strain in Experiment 11. **(g)** Relationship between the fraction of all amino acids and the sensitivity of growth to a ten-fold change in spatial separation from 25 $\mu$m to 250 $\mu$m (Experiments 5, 9-11 **Table 1**). The sensitivity of growth over distance, or interaction strength, is defined as the ratio of the minimum doubling times of a strain 25 $\mu$m to 250 $\mu$m from the partner strain ($D_{25}D_{250}^{-1}$). A positive interaction was defined as a statistically significant $D_{25}D_{250}^{-1}$ greater than one. A negative interaction was defined as a statistically significant $D_{25}D_{250}^{-1}$ less than one. **(h)** Relationship between the fraction of all amino acids and the sensitivity of growth of $\Delta pheA$ (interaction strength) to a ten-fold change in spatial separation from 25 $\mu$m to 250 $\mu$m from $\Delta metA$ (Experiments 5, 9-11 **Table 1**).

**Experiment 1 - Sender-receiver (step-response)**

| sender | 50 | 100 | 250 |
|---|---|---|---|
| 25 | 0.747801 | 0.022608 | 0.0845027 |
| 50 | | 0.184794 | 0.2742883 |
| 100 | | | 0.9255316 |

| receiver | 50 | 100 | 250 |
|---|---|---|---|
| 25 | 0.359228 | 0.000996 | 3.43E-07 |
| 50 | | 0.019142 | 6.91E-06 |
| 100 | | | 1.02E-06 |

**Experiment 2 - Sender-receiver (120 min oscillation) SNR, Fig. S6d**

| sender | 50 | 100 | 250 |
|---|---|---|---|
| 25 | 0.4142 | 0.4865 | 0.4724 |
| 50 | | 0.486 | 0.496 |
| 100 | | | 0.4009 |

| receiver | 50 | 100 | 250 |
|---|---|---|---|
| 25 | 0.0084 | 0.0946 | 0.0386 |
| 50 | | 0.1281 | 0.0038 |
| 100 | | | 0.3634 |

**Experiment 3 - Sender-receiver (60 min oscillation) SNR, Fig. 4i**

| sender | 50 | 100 | 250 |
|---|---|---|---|
| 25 | 0.3011 | 0.5635 | 0.6508 |
| 50 | | 0.2185 | 0.4167 |
| 100 | | | 0.5957 |

| receiver | 50 | 100 | 250 |
|---|---|---|---|
| 25 | 0.041243 | 0.000008 | 0.000063 |
| 50 | | 0.006158 | 0.007676 |
| 100 | | | 0.111161 |

**Experiment 4 - Dual-feedback oscillator**

**Fig. 4c, amplitudes**

| activator | 50 | 100 | 250 |
|---|---|---|---|
| 25 | 0.00027 | 0.000882 | 0.0003684 |
| 50 | | 0.989946 | 0.2593051 |
| 100 | | | 0.3396577 |

| repressor | 50 | 100 | 250 |
|---|---|---|---|
| 25 | 0.490548 | 0.00281 | 0.006595 |
| 50 | | 0.01196 | 0.026273 |
| 100 | | | 0.82516 |

**Fig. 4d, number of peaks**

| activator | 50 | 100 | 250 |
|---|---|---|---|
| 25 | 0.530488 | 0.082501 | 0.0121141 |
| 50 | | 0.074239 | 0.0180749 |
| 100 | | | 0.7831872 |

| repressor | 50 | 100 | 250 |
|---|---|---|---|
| 25 | 0.482493 | 0.116482 | 7.36E-06 |
| 50 | | 0.221686 | 1.12E-05 |
| 100 | | | 0.003286 |

**Fig. 4e, maximum correlation**

| | 50 | 100 | 250 |
|---|---|---|---|
| 25 | 0.3311 | 0.0868 | 0.0002 |
| 50 | | 0.2526 | 0.0003 |
| 100 | | | 0.009 |

**Fig. S9c, time lag**

| | 50 | 100 | 250 |
|---|---|---|---|
| 25 | 0.893 | 0.3204 | 0.0216 |
| 50 | | 0.1789 | 0.0149 |
| 100 | | | 0.1008 |

Figure S15: **P-values for quorum-sensing experiments.** Unpaired t-tests or bootstrapped hypothesis testing (**Materials and Methods**) were used to compute the p-values for Experiments 1-4 (**Table 1**). Highlighted values indicate statistically significant differences ($P < 0.05$).

Experiment 5 - Auxotroph (control), Fig. 5b,c

| phe | 50 | 100 | 250 |
|---|---|---|---|
| 25 | 0.108647 | 0.801381 | 0.4505158 |
| 50 | | 0.191799 | 0.1860261 |
| 100 | | | 0.6497227 |

| met | 50 | 100 | 250 |
|---|---|---|---|
| 25 | 0.968962 | 0.222213 | 0.667792 |
| 50 | | 0.175572 | 0.621371 |
| 100 | | | 0.43086 |

Experiment 6 - Auxotroph (coupled - 0X M, 0X F), Fig. 5 b,c

| phe | 50 | 100 | 250 |
|---|---|---|---|
| 25 | 0.364665 | 5.57E-06 | 6.29E-06 |
| 50 | | 1.05E-05 | 1.49E-05 |
| 100 | | | 0.0271837 |

| met | 50 | 100 | 250 |
|---|---|---|---|
| 25 | 0.070324 | 0.131259 | 0.288765 |
| 50 | | 0.750636 | 0.413298 |
| 100 | | | 0.622076 |

Experiment 8 - Auxotroph (rescue - Δmet), Fig. 5f

| phe GP1 | 50 | 100 | 250 |
|---|---|---|---|
| 25 | 0.489943 | 0.55232 | 0.8350058 |
| 50 | | 0.138334 | 0.5632902 |
| 100 | | | 0.3307124 |

| met | 50 | 100 | 250 |
|---|---|---|---|
| 25 | 0.896383 | 0.424424 | 0.774661 |
| 50 | | 0.394931 | 0.817224 |
| 100 | | | 0.535158 |

| phe GP2 | 50 | 100 | 250 |
|---|---|---|---|
| 25 | 0.182705 | 0.000366 | 0.0045164 |
| 50 | | 0.020797 | 0.0899014 |
| 100 | | | 0.6249524 |

M and F assays, Fig. 5g

| M, del-phe | 0.01X | 0.025X | 0.05X | 1.0X |
|---|---|---|---|---|
| 0X | 0.154073 | 0.162219 | 0.1289866 | 0.1045635 |
| 0.01X | | 0.697201 | 0.2347936 | 0.0101755 |
| 0.025X | | | 0.377097 | 0.1538273 |
| 0.05X | | | | 0.110678 |

| F, del-met | 0.01X | 0.025X | 0.05X | 1.0X |
|---|---|---|---|---|
| 0X | 0.118566 | 0.051864 | 0.031624 | 0.010126 |
| 0.01X | | 0.523527 | 0.119889 | 0.01876512 |
| 0.025X | | | 0.04259 | 0.00445829 |
| 0.05X | | | | 0.00081317 |

Experiment 9 - Auxotroph (coupled - 0.003X AAs), Fig. S13

| phe | 50 | 100 | 250 |
|---|---|---|---|
| 25 | 0.064692 | 0.000759 | 0.0003889 |
| 50 | | 0.120851 | 0.043729 |
| 100 | | | 0.2873524 |

| met | 50 | 100 | 250 |
|---|---|---|---|
| 25 | 0.037385 | 0.641391 | 0.55353 |
| 50 | | 0.098395 | 0.283497 |
| 100 | | | 0.803795 |

Experiment 10 - Auxotroph (coupled - 0.005X AAs), Fig. S13

| phe | 50 | 100 | 250 |
|---|---|---|---|
| 25 | 0.011602 | 0.009112 | 0.4962415 |
| 50 | | 0.673519 | 0.0291812 |
| 100 | | | 0.022791 |

| met | 50 | 100 | 250 |
|---|---|---|---|
| 25 | 0.247728 | 0.754728 | 0.082218 |
| 50 | | 0.505827 | 0.225389 |
| 100 | | | 0.147733 |

Experiment 11 - Auxotroph (coupled - 0.02X AAs), Fig. S13

| phe | 50 | 100 | 250 |
|---|---|---|---|
| 25 | 0.58644 | 0.040782 | 0.1821195 |
| 50 | | 0.288083 | 0.6915116 |
| 100 | | | 0.3050522 |

| met | 50 | 100 | 250 |
|---|---|---|---|
| 25 | 0.05376 | 0.943291 | 0.62213 |
| 50 | | 0.057607 | 0.163257 |
| 100 | | | 0.668231 |

Figure S16: **P-values for auxotroph experiments.** Unpaired t-tests or bootstrapped hypothesis testing (**Materials and Methods**) were used to compute the p-values for Experiments 5-11 (**Table 1**). Highlighted values indicate statistically significant differences ($P < 0.05$).
